# Exploring the Impact of Volumetric Additive Manufacturing of Photo-crosslinkable Gelatin on Mesenchymal Stromal Cell Behavior and Differentiation

**DOI:** 10.1101/2025.02.17.638591

**Authors:** Nele Pien, Bryan Bogaert, Marguerite Meeremans, Cezar-Stefan Popovici, Peter Dubruel, Catharina De Schauwer, Sandra Van Vlierberghe

**Affiliations:** Polymer Chemistry and Biomaterials Group, Centre of Macromolecular Chemistry (CMaC), Department of Organic and Macromolecular Chemistry, Ghent University, Krijgslaan 281 building S4, 9000 Ghent, Belgium; Veterinary Stem Cell Research Unit, Department of Translational Physiology, Infectiology and Public Health, Faculty of Veterinary Medicine, Ghent University, Salisburylaan 133, 9280 Merelbeke, Belgium

**Keywords:** tomographic volumetric additive manufacturing, photo-crosslinkable gelatin, bioresin development, mesenchymal stromal cells, MSC differentiation, cell-biomaterial interactions

## Abstract

This study investigates photo-crosslinkable gelatin-based hydrogels - thiolated gelatin (GelSH) and gelatin norbornene (GelNB) - for volumetric additive manufacturing (VAM). GelSH was synthesized with degrees of thiol substitution (DS) of 39%, 54%, and 63%, and GelNB with a DS of 60% (with respect to primary amine content). These were combined into GelNB-GelSH photo-resins at 5, 7.5, and 10% (w/v) and crosslinked via thiol-ene chemistry. Physico-chemical analysis showed that increasing DS and polymer concentration reduced swelling and increased moduli. VAM enabled the fabrication of high-resolution 3D hydrogel constructs from optimized formulations, demonstrating the ability to encapsulate mesenchymal stromal cells (MSCs) within a mechanically tunable, cell-supportive hydrogel environment. Film-cast hydrogels, also with embedded MSCs, served as comparative controls. VAM-printed constructs exhibited significantly higher alkaline phosphatase activity and calcium deposition, indicating enhanced osteogenesis. In contrast, chondrogenic and adipogenic differentiation were more pronounced in film-cast samples, due to their lower crosslinking density and stiffness. These findings emphasize the importance of matrix mechanics in guiding stem cell differentiation and demonstrate the potential of VAM for producing complex, functional scaffolds for tissue engineering. This work supports further development of tunable gelatin-based bioresins for applications requiring lineage-specific differentiation, including those targeting softer tissue types.

**Highlights:**

- Volumetric additive manufacturing (VAM) enables high-fidelity 3D hydrogel scaffolds.
- GelNB-GelSH bioresins support MSC encapsulation and differentiation.
- VAM scaffolds enhance osteogenesis, film-cast gels favor chondro- and adipogenesis.
- Mechanical properties and crosslinking density regulate stem cell fate in hydrogels.
- This study advances bioresin development for multi-lineage tissue engineering.

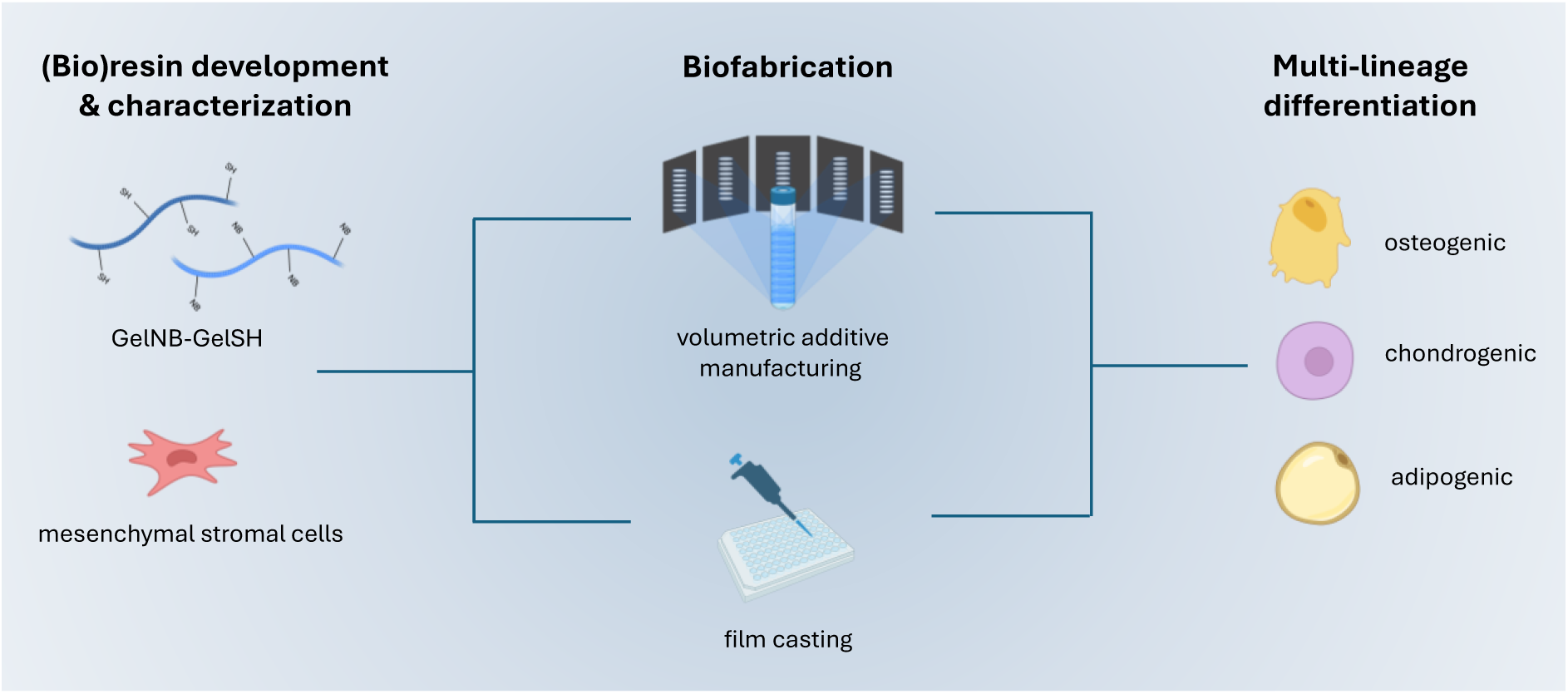

## 1. Introduction

**Volumetric additive manufacturing (VAM)** is a young 3D printing technique, pioneered by Kelly *et al.* [1], that represents a significant advancement over traditional layer-by-layer methods.[2] Unlike deposition-based 3D printing, which is limited by the sequential addition of layers, VAM utilizes light to polymerize a material throughout an entire volume simultaneously, enabling the rapid production of highly detailed, intricate and complex 3D structures in one single step.[2–4] Several volumetric light-based techniques have been developed showing unprecedented short printing times (down to a few tens of seconds) [5], while achieving good resolution (down to tens of micrometers, i.e., 20 µm) [6] and accuracy, making it ideal for creating scaffolds that mimic the architecture and function of native tissues.[7,8] The use of light-based polymerization in VAM also allows for fine control over the curing process, which is essential for creating well-defined scaffold features that promote cellular infiltration and tissue integration.[7,9] VAM has the ability to produce centimeter-scale, cell-laden constructs with minimal shear stress and photo-initiator exposure.

**Hydrogels** are particularly attractive for tissue engineering (TE) due to their biocompatibility, high water content, and ability to mimic the extracellular matrix (ECM) of tissues.[10] Gelatin, a biopolymer extracted from collagen, is an excellent choice for biofabrication due to its inherent bioactivity, biodegradability, and ability to support cell adhesion and growth. However, the incorporation of photocrosslinkable functionalities onto hydrogel building blocks is critical to enable precise control over their light-induced gelation process and mechanical properties during fabrication.[11] The foundational work by Kelly *et al.* [1] first demonstrated the applicability of VAM with 10 wt % gelatin-methacryloyl (**GelMA**) hydrogels (2 mM Ru/20 mM SPS), achieving ∼300 µm resolution at exposure doses above 100 mJ/cm². Volumetric printing of GelMA was further exploited by Buchholz *et al.* [12] to develop biofabricated human epithelial mammary ducts and endothelial constructs. In recent years, **thiol-ene click-like chemistries** and norbornene-based photo-resins have emerged as powerful alternatives to methacrylates.[13,14] This reaction is highly efficient, occurs under mild conditions, and allows for rapid crosslinking under light exposure (in the presence of a suitable photo- initiator).[15–17] The fast crosslinking kinetics and limited oxygen inhibition offered by the thiol-ene reaction are especially beneficial for VAM, where quick and uniform polymerization is required to achieve high-resolution prints. [18,19] Recently, Rizzo *et al.* [20] optimized a gelatin-norbornene (GelNB)-PEG4SH thiol-ene resin for fast (≈ 10-11s) volumetric prints supporting human dermal fibroblasts viability across 7 days of culture and C2C12 myotube formation; and later Chansoria *et al.* [21] produced multi-material auxetic lattices based on thiol-ene photoclick chemistry. In the work of Soliman *et al.* [22], GelNB hydrogels were employed to biofabricate complex architectures and to investigate vascularization through formation of microcapillaries, and Falandt & Levato achieved spatially selective grafting of thiolated growth factors in GelNB constructs to direct endothelial adhesion [23].

In TE, selecting appropriate materials and processing techniques is crucial, as these factors directly impact the scaffold’s biological and mechanical properties. The scaffold’s mechanical properties, such as stiffness and elasticity, must be tailored to meet the specific needs of the targeted tissue.[24,25] Likewise, the material’s ability to support cellular interactions and to guide cell behavior is equally important.[26,27] These material-processing interactions are particularly crucial when working with stem cells, such as **mesenchymal stromal cells (MSCs)**, which are capable of differentiating into a variety of tissue types, including bone, cartilage, and adipose tissue.[28,29] MSCs respond to mechanical cues and scaffold architecture to direct their differentiation. [24,30] Prior research has demonstrated that MSCs tend to differentiate into osteoblasts in stiffer substrates [31,32], i.e. an E-modulus between 11 and 30 kPa has been shown to promote osteogenic differentiation [33], whereas softer substrates promote adipogenic differentiation (E = 2 kPa) [34]. This illustrates that it is essential to carefully design scaffolds with the appropriate combination of material properties and processing techniques.[35–37]

Building on this, several groups have applied **VAM to MSC-laden constructs**: Bernal *et al.* [38] printed MSC-laden GelMA scaffolds for a trabecular bone model, showing osteogenic priming and endothelial co-culture, and Gehlen *et al.* [39] developed a perfusable vascularization model in GelMA with MSC/HUVEC co-printing and osteocytic marker expression over 42 days. Riffe & Burdick extended VAM to multi-material PEG/HA constructs embedding MSCs via a gelatin sacrificial network [40], while Kaplan & Zhang introduced pristine silk fibroin/sericin bioinks crosslinked through di-tyrosine by Ru/SPS to support hMSC osteogenesis of cell-seeded constructs [41]. Recent comparative work by Parmentier *et al.* [33] showed that thiol-ene-based inks require approximately a 3-fold less photo- initiator and drive MSC mineralization a 7-fold higher versus GelMA in both extrusion and VAM, and Duquesne *et al.* [4] reported a 2-fold increase in photo-crosslinkable moiety conversion and a 3-fold increase in bulk stiffness of the construct when comparing thiol-ene to GelMA in tomographic VAM, with concomitant increase in alkaline phosphatase activity. Together, these studies demonstrate VAM’s rapid maturation across diverse hydrogel chemistries and biological contexts. Some recent comprehensive reviews discuss these and other volumetric bioprinting breakthroughs, underscoring the field’s rapid evolution.[42,43]

Despite the diversity of VAM photo-resins and constructs, no study has directly compared VAM- printed versus film-cast GelNB-GelSH scaffolds to understand how fabrication-induced differences in crosslink density, stiffness, and microarchitecture influence MSC behavior. Here, we present the first side-by-side analysis of GelNB-GelSH63 (10 %(w/v)) hydrogels processed through volumetric printing and via static film-casting. We characterize their swelling, mechanical properties, and network crosslink density, then correlate these parameters with MSC viability, proliferation, and lineage-specific differentiation (osteo-, chondro-, adipogenic) over 21 days.

The aim of this study is to investigate how VAM biofabrication of GelNB-GelSH cell-laden hydrogels influences MSC proliferation and multi-lineage differentiation, providing insights into stem cell fate and advancing the use of VAM to serve tissue engineering applications.

## 2. Materials and methods

### 2.1 Materials

1-(3-Dimethylaminopropyl)-3-ethylcarbodiimide hydrochloride (EDC•HCl, TCI Europe, D1601, 25952- 53-8, Zwijndrecht); 2-mercaptoethanol (TCI Europe, M0058, 60-24-2, Zwijndrecht), 3-(4,5- dimethylthiazol-2-yl)-5-(3-carboxymethoxyphenyl)-2-(4-sulfophenyl)-2H-tetrazolium (MTS, Abcam, ab197010, 138169-43-4, Cambridge); 4-nitrophenyl phosphate (pNPP, ThermoFisher, 455010100, 330-13-2, Pittsburgh); 5-norbornene-2-carboxylic acid (Merck, 446440, 120-74-1, Darmstadt); (alcian blue, Merck, A3157, 33864-99-2, Darmstadt); alkaline phosphatase (ALP, Sigma, P7640, 9001-78-9, Darmstadt); boric acid (H3BO3, Merck, B6768, 10043-35-3, Darmstadt); calcein acetoxymethyl ester (Ca-AM, Merck, 56496, 148504-34-1, Darmstadt); calcium chloride (CaCl2, Sigma, C1016, 10043-52-4, Darmstadt); collagenase (ThermoFisher, 17100017, 9001-12-1, Pittsburgh); cresolphthalein (Sentinel Diagnostics, 17667H, 596-27-0, Milan); deuterium oxide (D2O, Eurisotop, D214H, 7789-20-0, Saint-Aubin); dialysis membrane (SpectraPor, MWCO 12-14 kDa, VWR, 734-0681, N/A, Pennsylvania); dimethyl sulfoxide (DMSO), Chem-Lab, CL00.0422.2500, 67-68-5, Zedelgem); ultrapure water (Milli-Q); Dulbecco’s phosphate buffered saline (DPBS, ThermoFisher, 14190250, N/A, Pittsburgh); ethanol (EtOH, Chem-Lab, CL00.0529.2500, 64-17-5, Zedelgem); ethylenediaminetetraacetic acid tetrasodium salt tetrahydrate (EDTA•4 H2O, Merck, 34103, 13235-36-4, Darmstadt); fetal bovine serum (FBS, Merck, F7524, 9014-81-7, Darmstadt); Fluorobrite^TM^ DMEM (Fluorobrite^TM^, ThermoFisher, A1896701, N/A, Pittsburgh); gelatin type B (Rousselot, , 9000-70-8, Ghent); guanidine•HCl (Carl Roth, 0037.3, 50- 01-1, Karlsruhe); hydrogen chloride (HCl, Merck, 258148, 7647-01-0, Darmstadt); iodixanol (OptiPrep^TM^, StemCell Technologies, 7820, 92339-11-2, Vancouver); isopropanol (IPA, Merck, 190764, 67-63-0, Darmstadt); L-glutamine solution (Merck, 25030024, 56-85-9, Darmstadt); low glucose Dulbecco’s modified eagle medium (Lg-DMEM, ThermoFisher, 11880028, N/A, Pittsburgh); N-acetyl- homocysteine thiolactone (Merck, A16602, 1195-16-0, Darmstadt); n-butylamine (Acros Organics, 219742500, 109-73-9, Geel); N-hydroxysuccinimide (NHS, Merck, 130672, 6066-82-6, Darmstadt); o-phthalaldehyde (OPA, Merck, P1378, 643-79-8, Darmstadt); Oil Red O (Merck, O0625, 1320-06-5, Darmstadt); Paraformaldehyde (Merck, 441244, 30525-89-4, Darmstadt); penicillin-streptomycin-amphotericin B suspension (ABAM, Merck, A5955, N/A, Darmstadt); phenylmethylsulfonyl fluoride (PMSF, Merck, 10837091001, 329-98-6, Darmstadt); phosphate buffered saline (PBS, Merck, P4417, N/A, Darmstadt); potassium chloride (KCl, Merck, 746436, 7447-40-7, Darmstadt); propidium iodide (PI, ThermoFisher, 15438249, 25535-16-4, Pittsburgh); PTFE-based release foil (ACC-3, Ventec International group); reverse osmosis water (Milli-RO); sodium bicarbonate (NaHCO3, TCI Europe, S0561, 144-55-8, Zwijndrecht); sodium carbonate (Na2CO3, TCI Europe, S0560, 497-19-8, Zwijndrecht); sodium hydroxide (NaOH, Merck, 221465, 1310-73-2, Darmstadt); tris(hydroxymethyl)aminomethane hydrochloride salt (TRIS•HCl, Merck, 10812846001, 1185-53-1, Darmstadt); triton X-100 (Merck, X100, 9036-19-5, Darmstadt); trypsin-ethylenediaminetetraacetic acid (Trypsin-EDTA, Merck, T4049, N/A, Darmstadt)

### 2.2 Development of photo-crosslinkable biomaterials

For the synthesis of thiolated gelatin (GelSH, Figure S1), a slightly adapted protocol from Meeremans *et al.* [44] was followed, employing three different molar ratios of amine to N-acetyl homocysteine thiolactone (1:1, 1:3, and 1:5) to achieve low, medium, and high degrees of substitution (DS) (Table S1). Gelatin-norbornene (GelNB, Figure S2) targeting a DS of 60%, was synthesized following a previously reported protocol [24]. The process involved activating 5-norbornene-2-carboxylic acid with EDC·HCl and NHS in DMSO, followed by conjugation to gelatin type B under inert conditions at 50°C for 16 hours (molar ratio of 1:0.75:1.5:1.2 for gelatin:EDC·HCl:NHS:5-norbornene-2-carboxylic acid, respectively). The product was purified through precipitation, dialysis and lyophilization to obtain the final GelNB.

### 2.3 Characterization of developed photo-crosslinkable materials

#### 2.3.1 ^1^H-NMR spectroscopy

The number of functional end groups in GelNB was determined using proton nuclear magnetic resonance (^1^H-NMR) spectroscopy [24]. In brief, 10 mg of the material was dissolved in 1 mL of deuterium oxide (D2O) and analyzed on a Bruker Avance WH 500 MHz NMR spectrometer at 40°C. The characteristic peaks of norbornene (6.0 and 6.3 ppm) were compared with the reference peak of the chemically inert hydrogen atoms of Valine (Val), Leucine (Leu), and Isoleucine (Ile) at 1.01 ppm. The NMR spectra were further analyzed using MestreNova software (version 6.0.2-5475), with baseline correction performed using the Whittaker smoother method. The following equations were used to calculate the DS for GelNB:

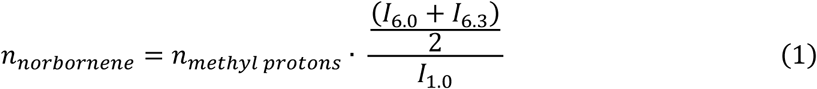

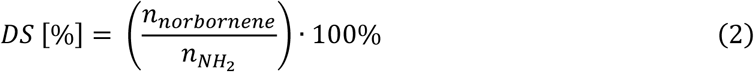

With:

Id = 6.3 - 6.0 ppm = integral of the signal of the protons of norbornene

Id = 1.0 ppm = integral of the signal of the protons of the reference peak

𝑛_𝑁𝐻2_ = 0.0385 moles primary amines per 100 g gelatin type B

𝑛_𝑚𝑒𝑡ℎ𝑦𝑙 𝑝𝑟𝑜𝑡𝑜𝑛𝑠_= 0.384 moles methyl protons (in reference signal) of Val, Leu and Ile per 100 g gelatin type B

#### 2.3.2 OPA assay to determine the degree of substitution of GelNB and GelSH

The number of functional end-groups in GelSH and GelNB was quantified using an o-phthalaldehyde (OPA) assay as previously described [45]. In brief, gelatin solutions were reacted with OPA and 2- mercaptoethanol, and the absorbance was measured at 335 nm via UV-VIS spectroscopy, with a blank for reference. A calibration curve with n-butylamine standards was used to calculate the degree of substitution based on the amount of unreacted amine groups (n = 3).

To enable consistent reference throughout the manuscript, we assigned specific names to the synthesized materials based on their DS, as determined via the OPA assay. Three derivatives of GelSH were prepared using 1, 3, and 5 molar equivalents of modifying agent (with respect to the primary amine content), resulting in DS values of 38.9 ± 4.5%, 53.8 ± 0.4%, and 62.8 ± 2.3%, respectively. These are referred to as **GelSH39**, **GelSH54**, and **GelSH63**, corresponding to their approximate DS percentages. For GelNB, a single formulation was synthesized using 1.2 equivalents with respect to primary amine content, with a targeted DS of 60%, and is referred to as **GelNB** throughout the manuscript.

#### 2.3.3 Preparation of solutions for photorheology and crosslinking of films

A stock solution of 0.8% (w/v) lithium phenyl-2,4,6-trimethylbenzoylphosphinate (LAP) was prepared by dissolving 40 mg of LAP in 20 mL of ultrapure water. An aqueous solution of GelSH (1, 3 or 5 eq) and GelNB was prepared using an equimolar ratio of thiol to alkene functionalities (GelNB-GelSH concentration of 5%, 7.5% and 10% (w/v)). Upon complete dissolution, 2 mol% (corresponding to 0.0076% (w/v)) of LAP was added with respect to the -NB functionalities. These mixtures were then used for comparative photorheology measurements and to prepare hydrogel films. An overview of the hydrogel formulations and their composition can be found in Supplementary Information, Table S2 (a).

#### 2.3.4 In situ crosslinking using photorheology

The UV-crosslinking behavior of the aforementioned materials was analyzed using photorheology with an Anton Paar Physica MCR 302e rheometer. The rheometer was equipped with an Omnicure S1500 light source, featuring a filter for wavelengths between 400–500 nm, which irradiated the sample (n = 3) from below through a quartz plate. A parallel plate setup with a 25 mm diameter top spindle was utilized. The curing process was monitored by deriving the storage modulus (G’) and loss modulus (G’’) of the solutions over time under light exposure (22.5 mW·cm^-2^) at 37°C, with a strain of 0.1% and a frequency of 1 Hz to ensure measurements were within the linear viscoelastic range. The initial gap between the plates was set to 0.300 mm, and the normal force was maintained at 0.5 N throughout the experiment. Measurements of the storage and loss moduli were taken before, during, and after light irradiation with interval durations of respectively 1 minute, 9 minutes, and 5 minutes.

#### 2.3.5 Photocuring of the solutions to prepare hydrogel films

The solutions (as described in section 2.3.3) were placed between two parallel glass plates that were covered with PTFE-based release foil and separated with a 1 mm thick silicone spacer. The material was then irradiated for 30 minutes from top and bottom with UV-A light (λ = 320-380 nm, 2 × 4 mW·cm^-2^) to ensure complete crosslinking. LAP has a local absorbance maximum at approximately 375 nm and significant absorbance at 365 nm (molar extinction coefficient at 365, ε=218 M^−1^cm^−1^).[46]

#### 2.3.6 HR-MAS ^1^H-NMR spectroscopy

The absolute crosslinking efficiency can be determined via high resolution magic angle spinning (HR-MAS) NMR spectroscopy [47]. Measurements were conducted on a Bruker Ascend™ 500 MHz spectrometer with a 4 mm ¹H/¹³C dual-channel HR-MAS probe at r.t. and a 6 kHz spinning rate. Freeze-dried samples were compressed in Kel-F inserts with 45 µL D₂O, sealed, and placed in the rotor. Quantitative data were obtained with a 30-second relaxation time, and the degree of conversion (DC) was calculated using the following equation:

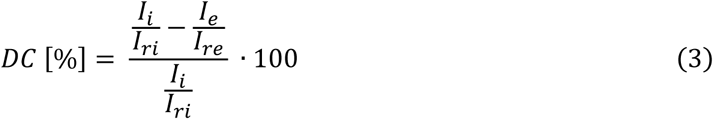

Ii = integral of the norbornene before crosslinking (6.0 – 6.3 ppm) Iri = integral of the reference peak before crosslinking (1 ppm)

Ie = integral of the norbornene after crosslinking (6.0 – 6.3 ppm) Ire = integral of the reference peak after crosslinking (1 ppm)

#### 2.3.7 Evaluation of the gel fraction and mass swelling ratio

Cylindrical samples (diameter: 6 mm, thickness: 1 mm) were punched out of the hydrogel films (see 2.2.5). The samples were lyophilized and weighed in dry state to determine the dry mass (*md1*). Next, the samples (n = 6) were incubated in ultrapure water at 37°C for 72 hours, the mass was recorded (*ms*) and the discs were lyophilized once more to determine the second dry mass (*md2*). The gel fraction and mass swelling ratio q were then calculated using the following equations:

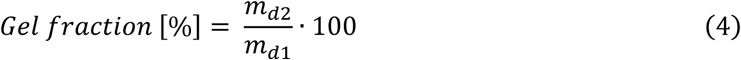

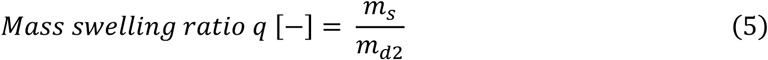

#### 2.3.8 Frequency sweep of crosslinked hydrogel samples

The viscoelastic properties of the materials were determined by a frequency sweep analysis. This was performed with a Physica MCR 301 rheometer from Anton Paar equipped with a parallel plate setup with diameter of 15 mm. First, hydrogel films were prepared according to section 2.3.5. The films were subsequently swollen to equilibrium in ultrapure water at room temperature for 72 hours. After reaching complete swelling, cylindrical samples of 14 mm (n = 3) were punched out. A frequency sweep analysis was performed from 0.1 to 10 Hz (FN = 0.5 N, amplitude of 0.1% and T = 37°C) to determine the storage (G’) and loss (G’’) moduli. The compressive modulus (E) was then calculated using the following equation:

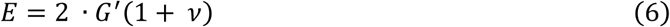

Where 𝜈 is the Poisson number assumed to be 0.5 for ideal hydrogels.[33]

#### 2.3.9 Tensile testing on crosslinked ring-shaped hydrogel samples

Hydrogel films were produced as described in section 2.3.5. Rings of 14 mm inner diameter and 18 mm outer diameter with a thickness ≈ 1.5 mm were punched out. The tensile properties of the equilibrium swollen GelNB-GelSH samples (n = 6) were determined at room temperature using a universal testing machine (Tinius Olsen 3ST) equipped with a 500 N load cell. The specimens were positioned as shown in Figure 1, between two 3D-printed curved hooks with a diameter of 5 mm. This method was developed specifically for gelatin-based hydrogels, instead of the traditional dogbone-shaped samples, that encounter issues when gelatin-based hydrogels are clamped between the grips.

**Figure 1.**
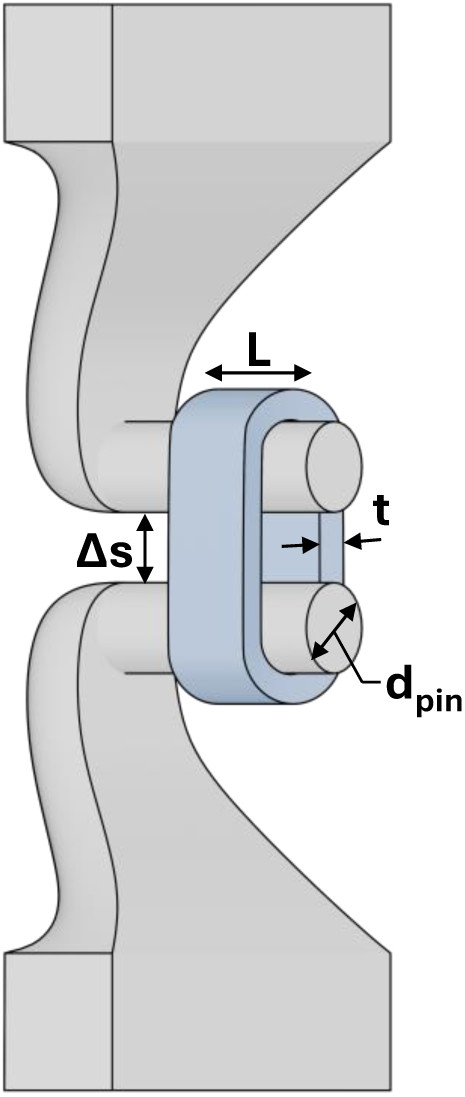
Tensile testing setup for hydrogel rings: Samples were mounted between 3D-printed curved hooks (dpin = 5 mm) and tested using a universal testing machine (n = 6). A preload force of 0.01 N was applied, and specimens were stretched at 5 mm·min^-1^. Displacement (Δs) was used to calculate the internal circumference (C), strain (ε), and circumferential stress (σ), where dpin is the hook diameter, L is the specimen length, and t is the specimen thickness. Young’s modulus was determined from the linear region of the stress-strain curve.

A preload force of 0.01 N was applied and the specimens were tested at a crosshead velocity of 5 mm·min^-1^. From this, the displacement and force were measured and transformed into stress-strain plots. Using the displacement, Δs, the internal circumference, C, was calculated using the following formula:

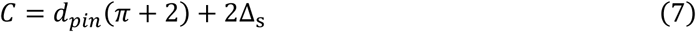

where dpin is the diameter of the 3D-printed hooks. The circumferential stress, σ, and the strain, ε, were calculated as the ratio of C with the initial internal circumference, Cinit, using the following formulas:

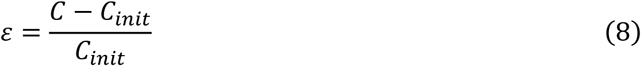

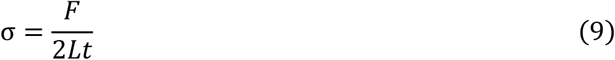

where F is the applied load during testing and L and t are the length and thickness of tubular specimen.

Young’s moduli were calculated from the initial linear region slope of the stress-strain plots.

### 2.4 Photo-resin preparation and characterization

#### 2.4.1 Preparation of the photo-resins

Aqueous solutions of GelSH5eq and GelNB in ultrapure water were prepared using an equimolar ratio of thiol to alkene. Upon complete dissolution, LAP stock solution was added to yield a final LAP concentration of 0.05% (w/v) and a final total gelatin concentration of 10% (w/v). An overview of the resin formulations and their composition can be found in Supplementary Information, Table S2 (b).

#### 2.4.2 Determination of gel point for dose optimization

Photorheology was performed to gain insight in the crosslinking kinetics as well as the optimal dose for volumetric printing. The photorheology analysis was performed as described in section 2.3.4 aside for a reduction of light intensity in order to increase the accuracy by delaying the cross-over point. The dose used as reference dose for volumetric printing (Dv) was calculated using the following formula:

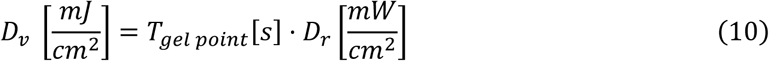

where Dr is the light dose used for photorheology and Tgel point is the time in seconds from the start of the light irradiation until the G’-G” crossover as shown in Figure S3.

#### 2.4.3 Absorption coefficient

The absorbance at 405 nm was measured using UV-VIS spectroscopy (UvikonXL, Bio-Tek Instruments) equipped with a temperature control unit. The temperature was set to 4°C and allowed to equilibrate for 30 minutes. The photo-resins were prepared as described in section 2.4.1. Subsequently, 500 µL was transferred to a PMMA micro-cuvette avoiding any entrapped air bubbles. The cuvette was inserted into the cooled spectrophotometer chamber and the absorbance of the sample was recorded after 7 minutes. The absorption coefficient 𝛼(𝜆) used for volumetric printing (Tomolite v1.0) was calculated using Equation 11, where 𝐴(𝜆) is the absorbance measured with the spectrophotometer, and t is the thickness of the sample:

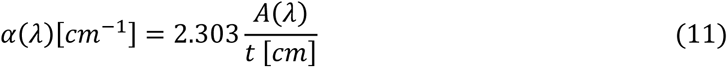

#### 2.4.4 Measuring the refractive index

The refractive index (RI) of the photo-bioresins were measured by applying 300 µL of the resin on the sensor of the refractometer (Deosdum). The refractometer was then placed in the fridge (4°C) for 7 minutes after which the RI was determined.

#### 2.4.5 Dose testing

The dose testing followed the Readily3D protocol. In brief, 3 mL of the resin was added to the cuvette, placed in a cuvette holder, stored in the fridge for 7 minutes and inserted into the Readily3D Tomolite v1.0 printer. Next, the resin was exposed to 405nm light at 25 different dose levels, by varying both irradiation time and light intensity. Each exposure was delivered in the form of an 800 μm diameter spot with a 1 mm gap between spots and a bottom margin of 4 mm. The diameters of the exposed spots were measured using optical light microscopy (Zeiss-Axiotech100HD/DIC) with ZEN Core software. The classification of the exposed spots was conducted based on four categories: "No dot" (invisible to the naked eye or under a microscope), "Barely visible" (if Dspot/Dexpected< 0.625 or visible to the naked eye), "Small" (if 0.75 > Dspot/Dexpected > 0.625), and "Large" (if Dspot/Dexpected > 0.75), where Dspot is the measured diameter [μm] and Dexpected is the set diameter [800 μm]. The optimized dose was determined as the last, fully printed isodose of this dose test. The optimized dose D was calculated using Equation 11.

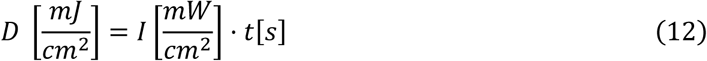

where *I* represents the intensity and *t* the illumination time.

### 2.5 Volumetric printing

#### 2.5.1 Scaffold design

Four different designs were VAM-printed (Figure 2), for various testing purposes. To determine the printing resolution of the photo-resins, a Schoen I-graph-wrapped package (IWP) was printed (Figure 2.A). Moreover, the CAD/CAM mimicry was evaluated on the benchmark from Figure 2.B, designed in Blender 4.0 graphical software with features ranging in sizes that challenge different aspects of printing resolution (positive vs negative resolution and z-axis vs x- and y-axis). For compression testing, cubes (dimensions: 5 mm x 5 mm x 5 mm) were printed, whereas discs (diameter of 8.8 mm and height of 1.5 mm) were printed for rheological measurements (frequency sweep analysis), as depicted in Figures 2.C and D respectively.

**Figure 2.**
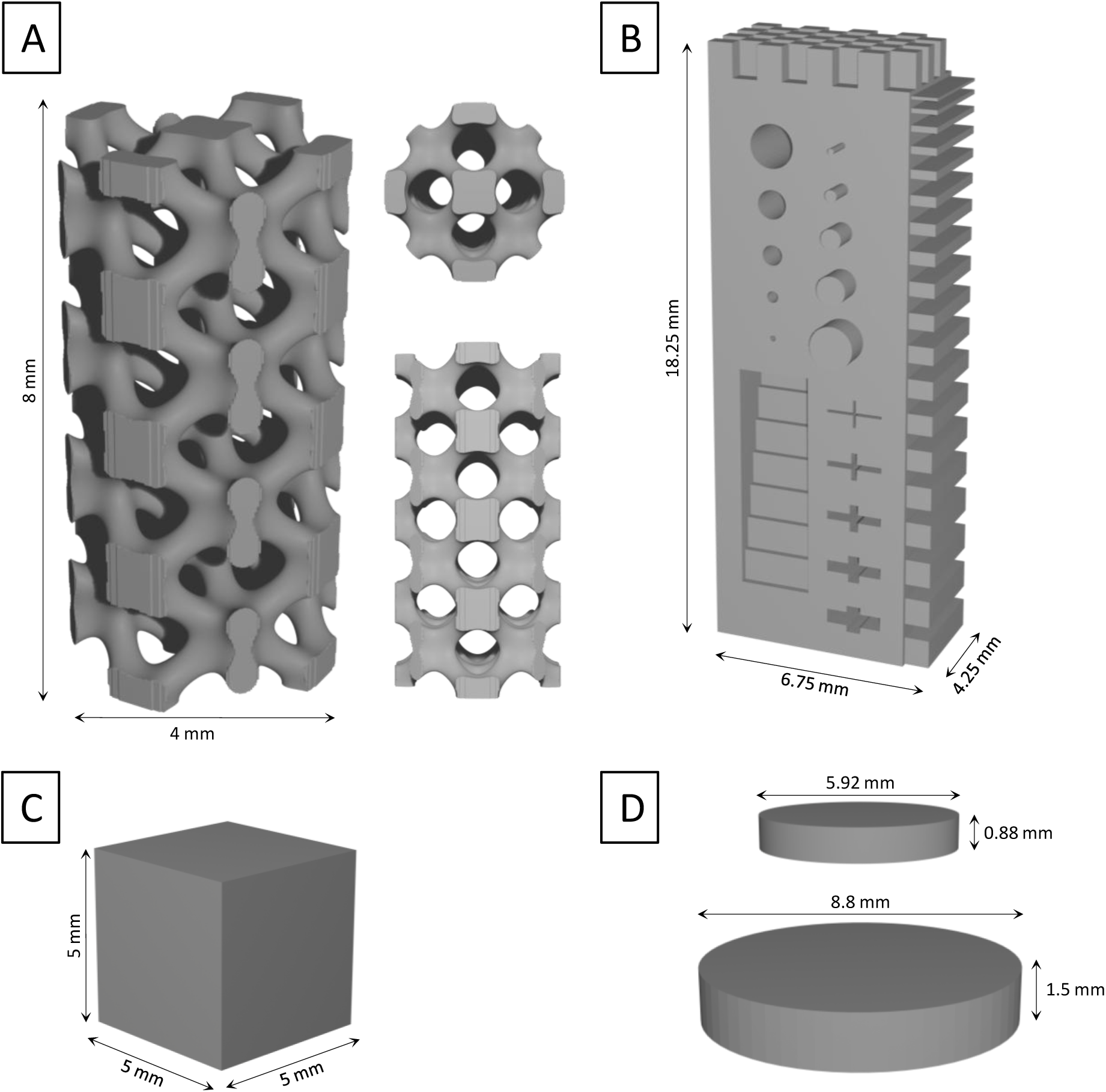
Designs that were volumetrically printed in this work for characterization of the scaffolds: (A) IWP design; (B) Benchmark 3D model for evaluation of the printing resolution and CAD/CAM mimicry; (C) Cubes (5x5x5 mm³) for compression testing; (D) Discs (diameter 8.8 mm and height 1.5 mm) for frequency sweep analysis (top) and discs with a diameter 5.25 mm and height 0.88 mm for cell encapsulation (bottom).

#### 2.5.2 Volumetric printing process

We used a commercially available Tomolite v1 system (Readily3D, 405nm light source), which implements the volumetric tomographic printing approach first reported by Kelly *et al.* [1]. No hardware modifications were made to the printer itself—our adaptations for gelatin-based hydrogels were carried out purely through photo-resin formulation (pre-print refrigeration, refractive-index matching, and dose calibration). Photo-resins were prepared according to section 2.4.1 and transferred to vials, ensuring no entrapped air bubbles were present. The vials were then placed in the fridge at 4°C for 7 minutes, before being transferred to the printing chamber of the volumetric printer. The STLs (depending on the purpose of the experiment) were loaded, and the parameters were set as follows; dose: 56 mJ•cm^-2^, refractive index: 1.334, voxel size: 25 µm. Once the printing process was complete, the vials were left in the printing chamber for 1 minute to allow for dark crosslinking.

#### 2.5.3 Post-printing processing

The vials were removed from the printing chamber and placed in a hot water bath (37°C) until the resin liquefied (30 - 90 seconds). The liquefied resin with printed structures was then poured into pre- heated PBS and washed for 2 minutes. Finally, the structures were transferred to a well plate, fresh PBS was added, and they were post-cured under UVA light or 405 nm light source (5, 10 or 15 minutes). The intensity of the light source was set at 8 mW·cm^-2^. Then, the double bond conversion percentage was assessed by HR-MAS NMR measurements (as described in section 2.3.6). A VAM-printed sample without post-processing curing was used as a negative control.

#### 2.5.4 CAD/CAM mimicry

The benchmark (Figure 2.B) was printed using the protocol from section 2.5.2. Optical light microscopy (Zeiss-Axiotech100HD/DIC) with ZEN Core software was used to measure the features immediately after the washing step. The dry state of the volumetrically printed scaffolds was visualized by scanning electron microscope (SEM, JCM-7000). Prior to SEM analysis, the samples were gold coated by sputter coating for 60 seconds at 15 mA under vacuum (K550X EmiTech).

#### 2.5.5 Mechanical characterization

a. *Compression testing on volumetrically printed cubes*

To conduct the compression test of the 3D printed hydrogels, cubes of 5x5x5 mm^3^ were VAM-printed using the protocol described in section 2.52 and 2.5.3. The cubes were then allowed to swell to equilibrium in ultrapure water for 72 hours at 37°C and were tested until failure at a compression rate of 2 mm·min^-1^ with a preload force of 0.01 N (Tinius Olsen 3ST) (n = 3).

b) Frequency sweep on volumetrically printed discs

To conduct oscillatory rheology of the VAM-printed hydrogels, discs with a diameter of 8.8 mm and a height of 1.5 mm were printed as described in section 2.5.2 and 2.5.3. The discs were allowed to swell to equilibrium in ultrapure water for 72 hours at 37°C and analyzed as described in section 2.3.8 (n = 3) using a parallel plate setup with a diameter of 8 mm.

### 2.6 Volumetric bioprinting with mesenchymal stromal cells

#### 2.6.1 Cell isolation and culture

Equine adipose tissue-derived MSCs were cultured under standard culture conditions, i.e. 38°C, 5% CO2, in expansion medium containing 20% FBS (Table 1). When a confluency of 80% was reached, cells were enzymatically detached with trypsin-EDTA (0.25% (w/v) trypsin, 0.53 mM EDTA) and counted. Cells of passage 4 were used for the experiments.

**Table 1.** Composition of media used for expansion (CTRL) and induction (DIFF).

| Expansion medium | Adipogenic Induction | Adipogenic Maintenance | Osteogenic | Chondrogenic |
| --- | --- | --- | --- | --- |
| DMEM-LG (Invitrogen, 11880-028) | DMEM-LG (Invitrogen, 11880-028) | DMEM-LG (Invitrogen, 11880-028) | DMEM-LG (Invitrogen, 11880-028) | Basal differentiation medium (Lonza, PT-3925) |
| 10% FBS | 15% rabbit serum (Sigma, R4505) | 15% rabbit serum (Sigma, R4505) | 10% FBS |  |
| - 1% = 2mM glutamine (Gibco, 25030024)<br>- 1% ABAM (Sigma, A5955) | - 1 μM dexamethasone (Sigma, D2915)<br>- 1% ABAM (Sigma, A5955) | - 10 μg·mL<sup>-1</sup> rh-insulin (Sigma, I9278)<br>- 1% ABAM (Sigma, A5955) | - 100 nM dexamethasone (Sigma, D2915)<br>- 10 mM β-glycerophosphate (Sigma, G9422)<br>- 0.05 mM L-ascorbic acid-2-phosphate (Fluka, 49752)<br>- 1% ABAM (Sigma, A5955) | - Single Quots differentiation medium (Lonza, PT-4121)<br>- 10 ng·mL<sup>-1</sup> Transforming Growth Factor-β3 (Sigma, T5425) |

#### 2.6.2 Preparation of the photo-bioresin

For volumetric printing with MSCs, a solution of the GelNB-GelSH5eq in DPBS was prepared. Iodixanol (OptiPrep^TM^, STEMCELL Technologies) stock solution (60%) was added to the GelNB-GelSH mixture to afford a final concentration of 0, 10, 15, or 20% (v/v) OptiPrep^TM^, to tune the refractive index (RI) of the bioresin to that of the cell cytoplasm to avoid undesired scattering and print defects. [48,49] Lastly, LAP stock solution (0.8% (w/v) LAP in DPBS) was added to yield a final concentration of gelatin of 10% (w/v) and a final LAP concentration of 0.05% (w/v). Upon complete dissolution, the solution was homogeneously mixed with MSCs (2 million cells·mL^-1^, passage 4). The photo-resins containing cells, further referred to as bioresins, were characterized by photorheology, UV-VIS spectroscopy, refractometry (see section 2.4).

#### 2.6.3 Parameters for volumetric bioprinting with MSCs

Two designs were VAM bioprinted: (1) disc (0.88 mm thickness, 5.28 mm diameter - which will result, after equilibrium swelling, into a disc with dimensions: 1 mm thickness and 6 mm diameter) and (2) the IWP structure (at 20% from the original .STL file). Volumetric printing was performed using the protocol from section 2.5.2. The post-processing curing consisted of 5 minutes under 405 nm light. The printed samples were then moved to a 96- or a 24-well plate, respectively. Expansion medium (100 µL or 1 mL) was added to each well, respectively. The medium was changed twice weekly.

### 2.7 Biological characterization of encapsulated MSCs

#### 2.7.1 Cell encapsulation in film-cast versus VAM-printed scaffolds

For the MSC encapsulation within a GelNB-GelSH hydrogel (later referred to as film-cast), and to evaluate the biological properties of the hydrogel itself: A 10% (w/v) GelNB-GelSH63 was prepared as described in section 2.3.3. Upon dissolution, the gelatin solution was homogeneously mixed with MSCs at a concentration of 2 million cells·mL^-1^. Then, 100 µL of this solution was pipetted in a 96-well plate. The cell-laden samples were placed in the fridge (10 min) for physical gelation, followed by 10 minutes of UV-A irradiation, as described earlier.[33] For the volumetric printing of the bioresin, and to evaluate the effect of the VAM printing process on the encapsulated MSCs, a GelNB-GelSH5eq (10% (w/v), 0.05% (w/v) LAP) solution was prepared in DPBS with 15% (v/v) OptiPrep^TM^. Then, 2 million MSCs·mL^-1^ were homogeneously mixed in and transferred to the VAM vial. The VAM printing was performed as described in section 2.6.3.

#### 2.7.2 Viability and proliferation assay

A 3-(4,5-dimethylthiazol-2-yl)-5-(3-carboxymethoxyphenyl)-2-(4-sulfophenyl)-2H-tetrazolium (MTS) assay was performed in order to assess the MSC metabolic activity. The expansion medium was removed and replaced with 120 µL MTS solution, i.e. 100 µL LG-DMEM and 20 µL MTS reagent. After an incubation period of 2 hours in the dark, the absorbance was measured using a plate reader (Multiskan SkyHigh Microplate Spectrophotometer, ThermoFisher Scientific) at 490 and 750 nm (as background). The MTS assay was performed on days 1, 7, 14 and 21 (n = 3). The hydrogel (without cells) was used to correct for background signal and results were normalized against Day 0 film-cast samples as a control group (100%).

A live/dead staining was performed using a dye solution of calcein acetoxymethyl (Ca-AM) and propidium iodide (PI) in Fluorobrite^TM^ DMEM to visualize the living and dead cells, respectively. The viability assay was also performed on days 1, 7, 14 and 21 (n = 3). The hydrogel (without encapsulated cells) was used as control group. On the evaluation days, the expansion medium was removed, and samples were washed twice with DPBS. Then, the dye solution containing 2 µL·mL^-1^ Ca-AM and 2 µL·mL^-1^ PI in Fluorobrite^TM^ DMEM was added to each sample (100 µL). After an incubation period of 15 minutes in the dark at room temperature, the dye solution was removed, the discs were washed with DBPS and Fluorobrite^TM^ DMEM was added. Then, (z-stack) imaging was performed using an inverted microscope (DMi1, Leica Biosystems). The raw data images were exported, and Fiji software version 2.9.0 was used to calculate the live/dead ratio.

#### 2.7.3 Preparation of media

The composition of the expansion medium (undifferentiated MSCs, CTRL) and the adipogenic, osteogenic and chondrogenic media (later referred to as differentiation media, DIFF) is provided in Table 1. To induce adipogenic differentiation, the encapsulated MSCs in GelNB-GelSH scaffolds were cultured in adipogenic induction medium for 72h, altered by 24h in adipogenic maintenance medium until analysis at day 21. For osteogenic and chondrogenic differentiation, the encapsulated MSCs were cultured up to 21 days.

#### 2.7.4 Differentiation assays

To assess alkaline phosphatase (ALP) activity, an early marker of osteogenesis, triplicate samples were washed with DPBS after 7 and 21 days of osteogenic differentiation and digested for 2 hours using 200 µL of collagenase suspension (1 mg·mL^-1^) per sample (n = 3). Following digestion, 150 µL of lysis buffer (consisting of 0.2% Triton X100 (Merck), 10 mM Tris•HCl at pH 8, and 0.5% phenylmethylsulfonyl fluoride (PMSF) in ultrapure water) was added. An 8-point standard curve was constructed by mixing 120 µL of p-nitrophenyl phosphate (pNPP) at varying concentrations (0–80 nmol·well^−1^) with 10 µL of ALP solution (1 mg·mL^-1^). Next, 80 µL of the diluted cell lysates were combined with 50 µL of pNPP solution. If necessary, further dilutions of the cell lysates were prepared to ensure the results fitted within the range of the standard curve. The absorbance was measured at 405 and 750 nm (Multiskan SkyHigh Microplate Spectrophotometer). ALP activity was then calculated as:

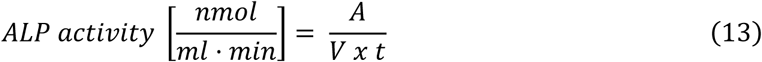

where A represents the amount of p-nitrophenol (pNP) produced (nmol), V is the volume of cell lysate added to the well (mL), and t is the reaction time (minutes). To enable comparison across all conditions, data were normalized to Day 7 film-cast samples cultured in expansion medium, which served as a control group (100%).

For calcium quantification, used as a late marker of osteogenesis, the samples (n = 3) were subjected to overnight digestion in 1 M HCl at 60°C (150 µL per sample) after 21 days of osteogenic differentiation. A standard curve for calcium detection was generated by serial dilutions of 1 M CaCl2 in ultrapure water to produce 8-point triplicate standards (0–1000 ng Ca^2+^ per well). Subsequently, 10 µL of either standards or diluted samples were mixed with 140 µL of a cresolphthalein-based working dye solution. After incubating the mixtures in the dark for 10 minutes at room temperature, the absorbance was recorded at 580 and 750 nm to measure the chromogenic purple complex formed between Ca^2+^ and cresolphthalein, with background correction applied as needed (Multiskan SkyHigh Microplate Spectrophotometer, ThermoFisher Scientific). To enable comparison across all conditions, data were normalized to film-cast samples cultured in expansion medium, which served as a control group (100%).

To assess chondrogenic differentiation of the encapsulated MSCs, Alcian Blue staining was used to detect glycosaminoglycans (GAGs). After removing the supernatant, cultures were rinsed twice with DPBS and fixed in 4% PFA for 2 hours at room temperature. Following fixation, samples (n = 3) were stained with 150 µL of 1% (w/v) Alcian Blue in 0.1 M HCl for 4 hours. Excess dye was washed away with PBS (4 times). For quantitative analysis, Alcian Blue was extracted by incubating the samples with 150 µL of 6 M guanidine HCl for 2 hours on a plate shaker. The extracted dye (50 µL) was mixed with an equal volume of 6 M guanidine HCl and measured using a spectrophotometer at 650/750 nm (Multiskan SkyHigh Microplate Spectrophotometer, ThermoFisher Scientific). Absorbance values were used to quantify GAG content, with blank wells serving as controls for background correction. Furthermore, data were normalized to film-cast samples cultured in expansion medium, which served as a control group (100%).

To assess adipogenic differentiation, Oil Red O staining was performed to detect lipid accumulation. Following removal of the medium, the cells were washed with DPBS and fixed with 4% PFA for 2 hours at room temperature (150 µL per well). After fixation, the wells were washed with ultrapure water. The cells were first incubated with 150 µL of 60% isopropanol per well for 5 minutes, followed by the addition of 150 µL of Oil Red O solution to each well for 15 minutes at room temperature with shaking. The Oil Red O solution was then removed, and the wells were briefly washed with 60% isopropanol for 30 seconds. To destain, 150 µL of 100% isopropanol was added to each well and incubated for 5 minutes with shaking. The resulting 100 µL of the destained solution was transferred to a new well, and absorbance was measured using a spectrophotometer at 510/750 nm (n = 3). The corrected absorbance was calculated by subtracting the OD 750 nm reading from the OD 510 nm reading to quantify lipid accumulation in the cells. For cross-condition comparison, all data were normalized relative to film-cast samples cultured in expansion medium, defined as the 100% reference standard.

### 2.8 Statistical analysis

Statistical analysis of the obtained data was conducted using Prism GraphPad Software version 10.2.2. The normality was assessed (using the Shapiro-Wilk test and by evaluating the Q-Q plots), and then a one-way or two-way ANOVA test with post-hoc multiple comparisons was applied to compare the different experimental groups. The selected tests are indicated in the captions of each figure. A p-value of less than 0.05 was considered statistically significant for differences between groups. The symbols representing the significantly different levels are indicated on the graphs, and/or defined in the captions (i.e. ns = p > 0.05; * = p ≤ 0.05; ** = p ≤ 0.01; *** = p ≤ 0.001; **** = p ≤ 0.0001).

## 3. Results and discussion

### 3.1 Material synthesis and characterization

GelSH and GelNB hydrogels were synthesized via thiol and norbornene functionalization respectively, with degrees of substitution (DS) quantified through the OPA assay and ^1^H-NMR spectroscopy (Supplementary Figure S4). Based on the OPA results, we refer to the thiolated gelatins as GelSH39, GelSH54, and GelSH63, corresponding to low, medium, and high DS levels, respectively. A single GelNB variant was synthesized with ∼60% DS. These functionalized components were combined at 5, 7.5, and 10% (v/v) to form GelNB-GelSH hydrogels.

Photorheology revealed tunable mechanical properties depending on DS and concentration, with storage moduli (G′) ranging from 2.1 ± 0.9 kPa for GelNB-GelSH39 at 5% (w/v) to 12.5 ± 1.8 kPa for GelNB-GelSH63 at 10% (w/v) (Figure S5). These values span the physiologically relevant stiffness ranges associated with adipogenic, chondrogenic and early osteogenic differentiation. All formulations showed high gel fractions (95–100%), indicating efficient crosslinking, with GelNB- GelSH63 at 10% showing the highest (99.7%). Mass swelling ratios ranged from 60 (GelNB-GelSH39 at 5%) to 20 (GelNB-GelSH63 at 10%), being inversely related to polymer concentration and crosslinking density (Figure S6). These trends highlight the capacity of the system to finely tune the hydrogel’s physicochemical and mechanical properties through simple adjustments in formulation. Full synthesis protocols, characterization data, and an extended discussion are provided in the Supplementary Information (Sections S1–S3, Figures S4–S6).

The mechanical characterization of GelNB-GelSH hydrogels through frequency sweep analysis and ring tensile testing provides valuable data on the hydrogels’ viscoelastic properties and tensile strength. These properties can be correlated with the biophysical cues that influence MSC differentiation.[31,33,50] For example, the storage modulus obtained from frequency sweeps can indicate the stiffness of the hydrogel, a critical factor in determining the differentiation pathway of encapsulated MSCs. However, tensile testing is particularly important for assessing the hydrogel’s ability to withstand macro-scale forces encountered *in vivo*, such as those associated with movement, muscle contraction, or blood flow.[37] Tensile testing provides information such as the ultimate tensile strength, the elongation at break, and the Young’s modulus, which are critical for predicting mechanical integrity and durability under physiological conditions.[51] These data can aid in determining whether the hydrogel will maintain its structure and support cellular activities, such as migration and adhesion, within a dynamic *in vivo* environment.

Given that the cells would be in a swollen equilibrium environment, mechanical properties were assessed via frequency sweep analysis (on equilibrium swollen hydrogel samples). The hydrogels were subjected to a constant 0.5N force under a rheometer spindle and underwent a frequency sweep. The mean storage and loss moduli (at 1 Hz) are presented in Figure S7. The compressive modulus E was calculated using Equation 6, assuming a Poisson’s ratio of 0.5.[33] As observed previously, decreasing concentration and functionalization degree led to lower mechanical strength, albeit not statistically significant. This trend can be attributed to the decrease in crosslinking density. GelNB-GelSH63 at 10% (w/v) exhibited the highest E at 10.0 ± 2.4 kPa, while GelNB-GelSH54 at 5% (w/v) displayed the lowest E at 3.2 ± 2.2 kPa. By tailoring the mechanical properties of the hydrogel, we aim to guide MSCs towards specific cell lineages. Considering that E-moduli around 2 kPa favor adipogenic differentiation [34] and 11-30 kPa promote osteogenesis [33], our hydrogels span a range relevant for various lineages.[48,52] The material with the highest E (GelNB-GelSH63 at 10% (w/v), E = 10.0 ± 2.4 kPa) shows promise for osteogenic differentiation, while the lower E values (e.g., GelNB-GelSH54 at 5% (w/v), E = 3.2 ± 2.2 kPa) may be more suitable for chondrogenic or even adipogenic differentiation. This suggests that by adjusting the hydrogel formulation, we can potentially cover the full range of mechanical properties required to guide differentiation towards osteogenic, chondrogenic and adipogenic lineages. This suggests that by adjusting the hydrogel formulation, we can potentially cover the full range of mechanical properties required to guide differentiation towards osteogenic, chondrogenic, and adipogenic lineages. Furthermore, the concentration of the hydrogel can influence other important factors, including the printing resolution and the porosity. Higher concentrations may lead to improved printability but can also result in reduced porosity, potentially affecting cell-material interactions and nutrient and waste product diffusion. Optimizing the hydrogel concentration is essential for achieving the desired balance between mechanical properties and structural characteristics.

In addition to the frequency sweep analysis, tensile testing on ring-shaped samples was also conducted. Ring-shaped samples were used for tensile testing instead of dogbone-shaped samples to prevent failure at the grips, as gelatin-based hydrogels are prone to tearing under the harsh clamping conditions needed to prevent slippage. The ultimate tensile strength (UTS), the Young’s modulus (YM), and the maximum strain were determined (Figure 3). The UTS, which measures the maximum stress a material can withstand before failure, showed a consistent trend, increasing with the polymer concentration. The highest UTS was observed in GelNB-GelSH63 at 10% (w/v), with a value of 23.88 ± 4.78 kPa. Notably, there was no significant difference in UTS when comparing the 10% (w/v) formulations for GelNB-GelSH39, 54 and 63 substitutions (i.e. 1, 3 and 5 eq). Similarly, the YM, reflecting the stiffness of the hydrogels, increased with the polymer concentration. The GelNB-GelSH formulations at 5, 7.5 and 10% (w/v) exhibited Young’s moduli of 0.6 ± 0.1, 0.7 ± 0.1, and 1.2 ± 0.2 kPa for the GelNB-GelSH63 variant, respectively. This is in agreement with a previous report from Yu *et al.* [52] who demonstrated that tensile moduli ranged between 0.5-5 kPa for GelNB-GelSH hydrogels with increasing concentrations between 5 and 15% (w/v) (molar ratio 1:1). No statistically significant difference in maximum strain was observed across the different formulations, despite the variations in UTS and YM. One potential explanation for this is that the crosslinking density, while impacting stiffness and strength, may have reached a plateau where further changes do not significantly affect the extensibility or strain at failure. Parmentier *et al.*[24] investigated this elastic behavior of gelatin hydrogels in detail and demonstrated that with increasing crosslinking densities of GelNB gels, no changes in elasticity could be observed. Additionally, the viscoelastic nature of the hydrogels might contribute to energy dissipation, resulting in similar strain values across formulations. It can thus be concluded that while higher concentrations and substitution degrees improve the mechanical strength and stiffness, the strain behavior remains consistent. As mechanical strength and stiffness are key factors for supporting osteogenic differentiation, which is rather difficult to reach with natural hydrogels, the GelNB-GelSH63 formulation at 10% (w/v) was selected for biofabrication using VAM and film casting.

**Figure 3.**
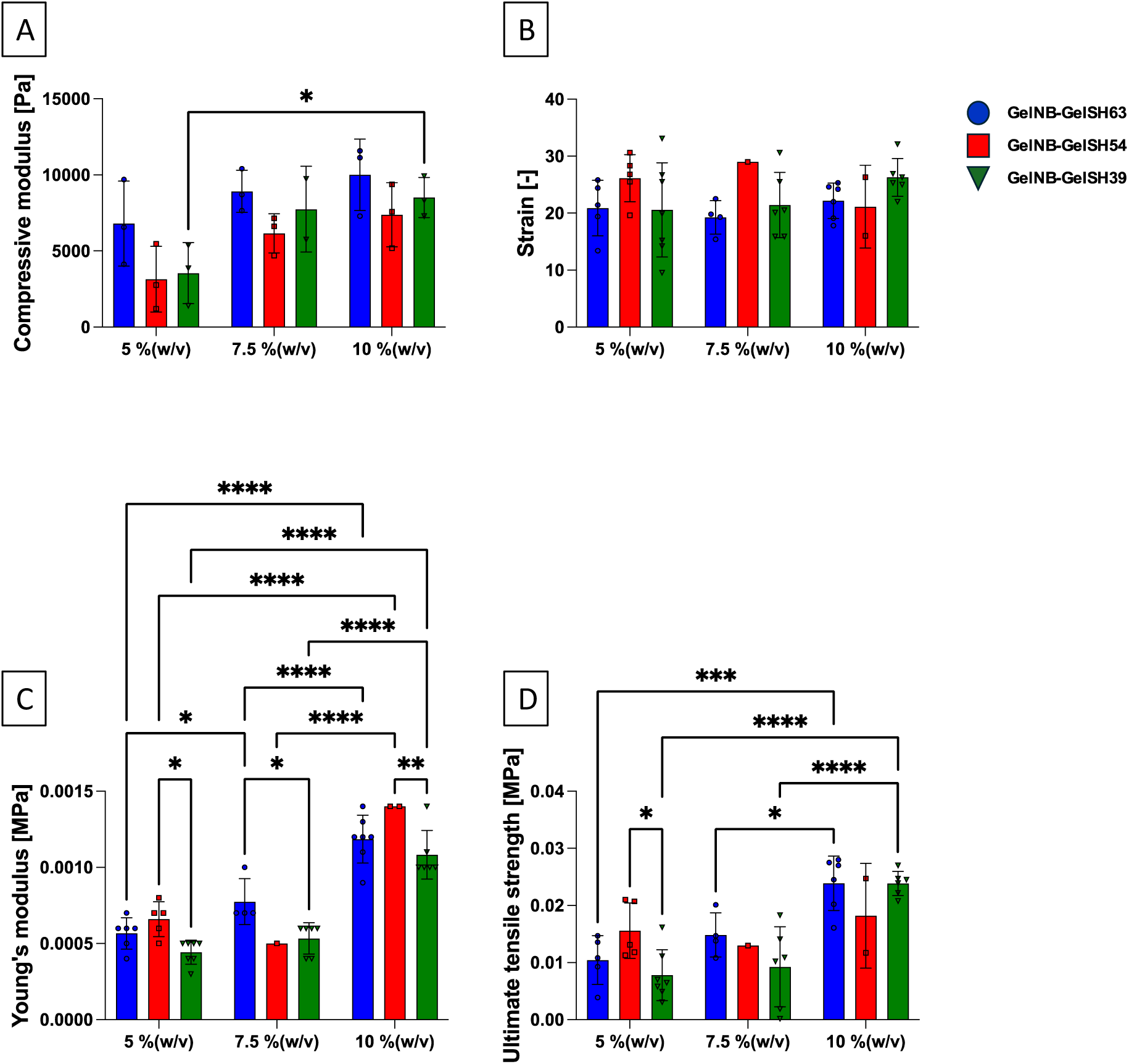
Mechanical properties of the crosslinked gelatin-based hydrogels (at equilibrium swelling): Mean compressive modulus determined via frequency sweep analysis on equilibrium swollen crosslinked discs (A); Strain [-] (B); Young’s modulus [MPa]; (C) and Ultimate tensile strength [MPa]; (D) determined via tensile testing on ring-shaped samples (n = 6). A two-way ANOVA and a Tukey’s multiple comparisons test was selected. (* = p ≤ 0.05; ** = p ≤ 0.01; *** = p ≤ 0.001; **** = p ≤ 0.0001)

### 3.2 Volumetric printing of GelNB-GelSH

#### 3.2.1 Resin development and characterization

First, an acellular photo-resin was developed to investigate the printability as well as the highest obtainable resolution of the GelNB-GelSH63 formulation. A higher% (w/v) of photo-initiator is needed in volumetric printing (as compared to the traditional film casting) because the entire 3D structure is cured simultaneously, requiring sufficient light absorption and uniform polymerization throughout the resin volume, even within complex scaffold geometries.[48,53,54] Based on reported values for similar hydrogel systems (0.05% (w/v) LAP for 5% (w/v) GelNB:4-arm-PEG-thiol [20], 0.05% (w/v) LAP for GelNB-GelSH [21] and 0.1-0.5% (w/v) LAP for 5% (w/v) GelMA [54,55]), and following optimization, a final concentration of 0.05% (w/v) LAP was selected for VAM printing in this work. Whereas oxygen acts as a radical scavenger for acrylate-based systems, in thiol-ene crosslinking an inhibitor (e.g. 2,2,6,6-tetramethyl-1-piperidinyloxy (TEMPO)) is typically added in order to obtain the necessary non-linear conversion response to light dose.[1,5] This is attributed to the reactivity difference of the thiyl radical addition to (strained) alkenes compared to the slower addition reaction to dissolved oxygen in contrast to the carbon centered radicals that react more easily with oxygen to form the less reactive peroxyl radicals.[56,57] On the other hand, in gelatin-based systems the natural anti-oxidant behavior of the amino acids (i.e. lysine, tyrosine and others) can act as an inhibitor and allows to obtain a non-linear response curve without the addition of dedicated radical scavengers like TEMPO. Next, the absorbance and the refractive index of the resin at 405 nm were determined to be 0.3738 and 1.334, respectively. Lastly, the optimal dose was determined by a combination of photorheology and the dose test. Using photorheology (Figure S8.A), a non-crosslinking interval of 5.5 seconds was observed, which allows sufficient light penetration without attaining the necessary dose for crosslinking, a crucial requirement for volumetric printing. In addition, the gel point, as obtained after plotting all data, was 6.2 seconds corresponding to a dose of 45 mJ·cm^-2^ which was used as the initial reference dose. To further refine this result under conditions more representative of the volumetric printing process, a second dose test was conducted following the Readily3D dose test (Figure S8.B). This test yielded an optimized dose of 56 mJ·cm^−2^, reflecting differences in light distribution and exposure dynamics between photorheology and the volumetric setup.

#### 3.2.2 Post-processing curing

In a next part of the work, we evaluated the curing efficiency of gelatin hydrogels comparing 405 nm with UV-A (320-380 nm) irradiation. The choice of wavelength is critical, especially for applications involving cell encapsulation, where minimizing cytotoxicity is essential. Light at a wavelength of 405 nm is generally preferred for its lower cytotoxicity, while UV-A offers more rapid curing.[58,59] Our focus was on determining the crosslinking efficiency through HR-MAS NMR spectroscopy while keeping irradiation times as short as possible to ensure high cell viability. The results (Figure S9) showed no significant differences in double bond conversion between the two light sources nor across the different curing times (5, 10, and 15 minutes). Without post-processing curing, the double bond conversion was 60%. After post-processing curing, high conversion rates (above 90%) were observed at all time points, demonstrating that the post-processing process was rapid and effective in achieving complete crosslinking. The absence of significant variation between the two light sources suggests that both UVA and 405 nm wavelengths are equally effective in promoting hydrogel formation. Therefore, either can be used for post-curing without compromising crosslinking quality. Importantly, 405 nm was identified as a preferable light source for crosslinking due to its reduced negative impact on cells [60,61], while still achieving a high double bond conversion, even at shorter irradiation times (96.2% at 5 minutes).

### 3.3 VAM-printed scaffold characterization

#### 3.3.1 CAD/CAM mimicry

The CAD/CAM mimicry achieved through volumetric printing is shown in Figure 4. The left panel presents an IWP structure with a lattice configuration and interconnected pores. The CAD model captures the porous architecture and overall cylindrical shape, which is faithfully reproduced in the printed scaffold. This highlights the capability of the VAM system to accurately and quickly translate complex 3D geometries into physical structures with high precision. The IWP design (Figure 4.A), characterized by its repeating porous architecture, serves as an example of how VAM can produce intricate, scaffold-like structures suitable for biomedical applications. The right panel showcases a more intricate benchmark CAD design (Figure 4.B), featuring circular holes, pillars, steps, crosses, and thin walls. The smallest observed features include holes and pillars at 340 µm, steps at 210 µm, and crosses and walls at 120 µm, as shown in the red markers on the CAD design (Figure S10). In comparison, values found in literature for gelatin polymer solutions include a positive resolution of 138.83 ± 52.47 μm and negative resolution of 198.97 ± 51.84 μm for GelNBNB+GelSH containing 0.025% (w/v) LAP [4], a negative resolution of ± 380 µm for GelNB+4-arm thiolated poly(ethylene glycol) containing 0.075% (w/v)% LAP [8], a positive resolution of 244 ± 16 µm and a negative resolution of 461 ± 26 µm for GelMA with 5% (w/v) LAP [62], as well as 23.68 ± 10.75 µm for GelNB+dithiothreitol/diethylene glycol with 0.1% (w/v) LAP [23]. The photographic image (Figure 4.C and 4.D) confirms the successful replication of these intricate details, with the VAM-printed structure retaining 109.01% of the original CAD dimensions after equilibrium swelling. Further evaluation using optical microscopy and SEM imaging (Figure S10 and S11) reveals that, although some challenges such as slight rounding of edges and difficulties in printing positive features were encountered, the majority of the features were accurately reproduced. SEM images of the printed benchmark (Figure S12) show walls with a thickness of 85 μm, closely matching the CAD design. This combination of optical and SEM imaging confirms the high fidelity of VAM printing in replicating complex CAD designs, with only minor deviations in the smallest, most intricate features.

**Figure 4.**
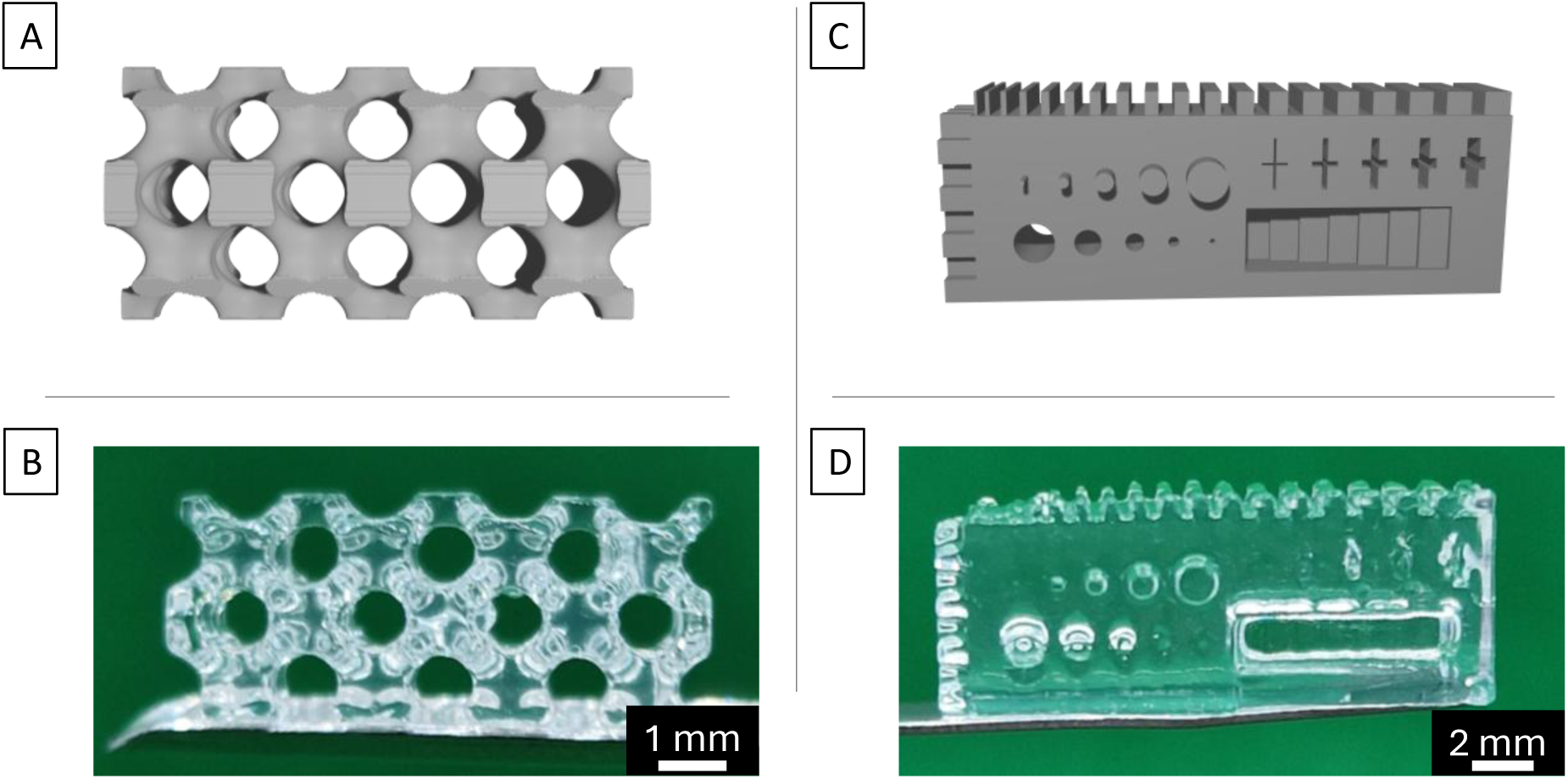
CAD/CAM mimicry of VAM-printed structures: (A) 3D image of the IWP design (CAD) and (B) the VAM-printed IWP design (CAM); (C) 3D image of the benchmark (CAD) and (D) the VAM-printed benchmark (CAM).

VAM printing outperforms conventional layer-by-layer methods in both fidelity and cell-friendly processing. Extrusion bioprinting of GelNB-GelSH hydrogels can achieve >90 % fidelity for ∼400–700 µm features but is limited by nozzle size, slow deposition and imparts shear stresses that can reduce cell viability or alter the cell phenotype [52]. Digital light processing offers sub-100 µm XY resolution, yet, in GelSH-GelNB resins, rapid surface thiol oxidation induces disulfide formation, which further reduces the printability of such resins. By contrast, VAM’s tomographic curing offers ∼20 µm resolution within tens of seconds, eliminates inter-layer artifacts, and avoids nozzle-induced shear.

#### 3.3.2 Mechanical characterization

The mechanical properties of VAM-printed hydrogel cubes composed of GelNB-GelSH63 at 10% (w/v) were evaluated through compression testing and oscillatory rheology, as outlined in section 2.5.5. After swelling, the cubes measured approximately 5.25, 5.25, 5.12 mm in length, width and height, respectively. The mechanical tests showed consistent behavior across the triplicate samples (Figure 5). The hydrogels initially exhibited a linear elastic response, where the force increased proportionally with the displacement. However, at higher forces, the elasticity became non-linear. As compression continued, the hydrogels eventually reached a critical stress point, leading to structural collapse. The hydrogels demonstrated a maximum strain of 69.7% and withstood a considerable compressive force before failure, with an average compressive strength of 507.63 ± 53.15 kPa.

**Figure 5.**
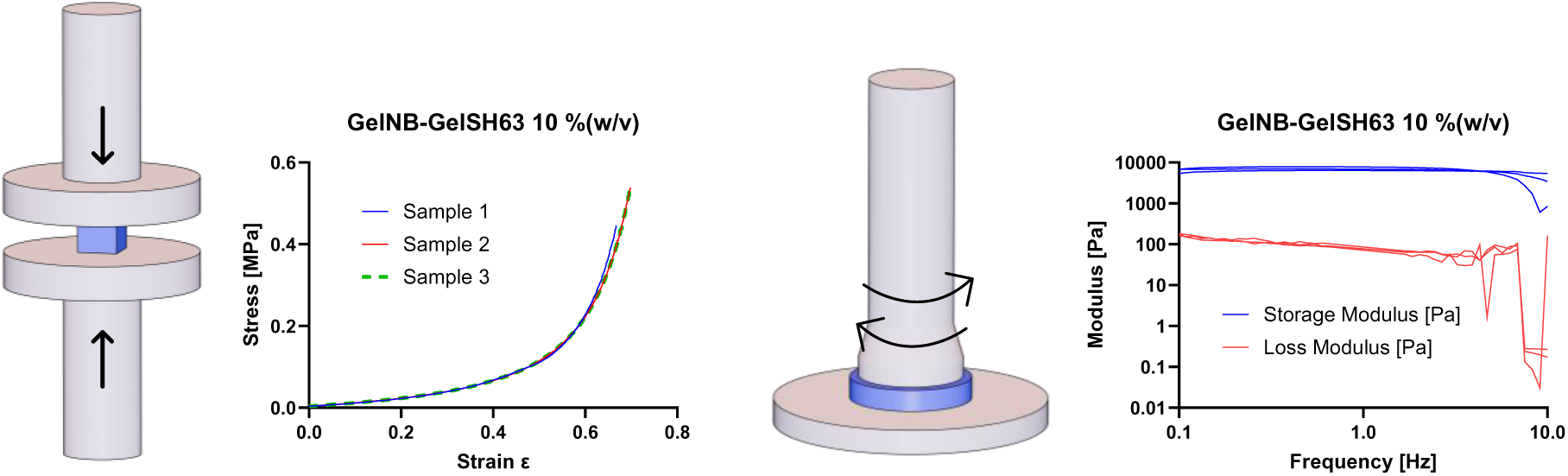
Compression testing and frequency sweep analysis on VAM-printed cubes and discs (n = 3), respectively.

VAM-printed GelNB-GelSH63 discs, with a diameter of 8.8 mm and a height of 1.5 mm after equilibrium swelling, were evaluated using oscillatory rheology. The analysis revealed that these hydrogel scaffolds exhibited a storage modulus of 7.014 ± 0.715 kPa and a loss modulus of 0.077 ± 0.007 kPa, reflecting their viscoelastic nature. This corresponds to a compressive modulus (E) of 21.04 ± 2.14 kPa. To compare, frequency sweep analysis of film-cast GelNB-GelSH63 samples (at 10% (w/v)) indicated a storage modulus of 2.156 ± 0.515 kPa, corresponding to an E modulus of 6.467 ± 0.154 kPa. This value is approximately 3.5 times lower than that of the VAM-printed scaffolds, suggesting significant differences in their mechanical properties. These differences can be attributed to variations in double bond conversion and as a result the mechanical properties, as demonstrated by HR-MAS (95.0% for VAM-printed *versus* 41.7% film-cast, Figure S13). This increased conversion is largely due to differences in processing methods. Specifically, the post-processing curing step in VAM printing enhanced double bond conversion from 60.6% to 95.0% (see section 3.2.2), whereas the cell encapsulation protocol for film-cast constructs involved 10 minutes of physical crosslinking followed by 10 minutes of UV irradiation, with a lower PI concentration.[33,63]

### 3.4 Optimization of the bioresin for volumetric bioprinting

In the previous section, an acellular photo-resin was prepared, volumetrically printed and characterized. However, to achieve the goal of encapsulating cells within the final scaffolds, further optimization steps were necessary. The inclusion of cells introduced light scattering effects, which can lead to print defects and reduced resolution. Additionally, the presence of cells affected the photopolymerization dynamics, prompting us to reassess and refine the photorheology to ensure consistent crosslinking and scaffold fidelity. An important aspect when performing VAM with cells is to tune the refractive index (RI) of the bioresin to that of the cell cytoplasm to avoid undesired scattering and print defects. The cytoplasm typically has a RI between 1.36 and 1.39, depending on the cell type.[48,64–66] Tuning of the RI can be achieved via addition of iodixanol, a cell-compatible, radio-opaque contrast agent with a refractive index of 1.429 (at 60% (w/v), commercially available as OptiPrep^TM^).[66–68] Maintaining an iodixanol concentration as low as possible is essential as iodixanol has also been shown to impact the ductility and overall mechanical performance of scaffolds.[38,55,69] In brief, bioresins were supplemented with 0, 10, 15 or 20% (v/v) OptiPrep as well as MSCs at a concentration of 2·10^6^·mL^-1^ while retaining a concentration of 10% (w/v) GelNB-GelSH63 and 0.05% (w/v) LAP. The lowest RI within the range for the cytoplasm was found using 15% (v/v), being 1.3623 (other RI can be found in Figure S14 and Table S3). To confirm that OptiPrep® addition did not significantly diminish mechanical performance, photo-rheological analysis was carried out on each formulation (Figure S15). The complete bioresin exhibited an absorbance of 0.587 at 405 nm. The gel point was determined using photorheology (Figure S16) and served as the initial reference dose (36 mJ·cm^-2^) for the subsequent second-dose test (Figure 6.B). The second dose test was then performed using the Readily3D protocol for VAM printing, yielding an optimized dose of 71 mJ·cm^-2^.

**Figure 6.**
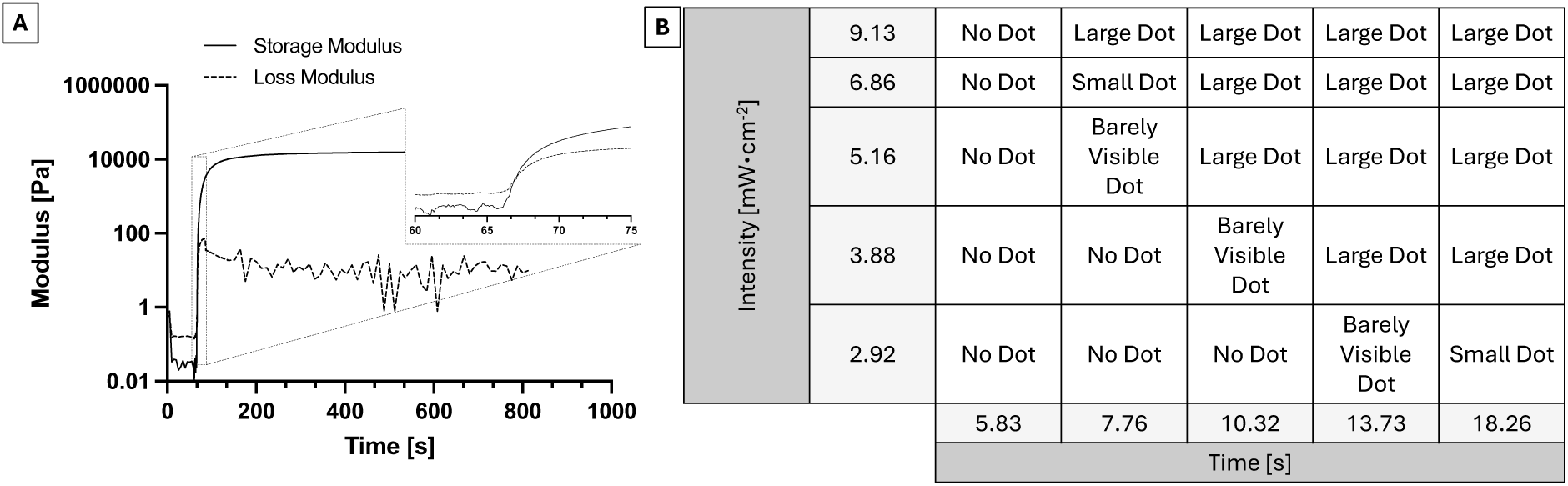
Determination of the reference dose for VAM printing. (A) Photo-rheology analysis to determine the gel point and required light dose for hydrogel crosslinking. The storage modulus (solid line) and loss modulus (dashed line) were monitored over time, with a gelation point observed at the crossover. The inset highlights the modulus increase during the gelation process. (B) Evaluation of reference doses using the Readily3D protocol, showing the relationship between light intensity (mW·cm^−2^) and exposure time (s) to achieve effective crosslinking.

### 3.5 Biological characterization of encapsulated MSCs

Recently, interest has grown in understanding how different microenvironments, such as those provided by hydrogels, influence the behavior of MSCs, particularly when comparing film-cast cells versus printed cells.[70–72] This study aimed to evaluate the biological properties, specifically proliferation and tri-lineage differentiation, of MSCs processed using two methods: biofabrication using VAM printing and encapsulation via film casting. By assessing these differences, we sought to determine how the structural variations introduced by these techniques impact cell behavior.

#### 3.5.1 Viability and proliferation assay

The MTS assay results, normalized to Day 0 film-cast samples, provide valuable insights into the proliferation of encapsulated MSCs during 21 days of culture. Both groups, VAM-printed versus film- cast, exhibited significant increases in absorbance, indicating successful proliferation (Figure 7.A), with a notable interaction between both the processing method and time points of the measured outcomes (*p < 0.0001*). On day 0, there was a significant difference between VAM-printed and film- cast samples (*p = 0.0027*), with film-cast samples showing higher absorbance values. This can be attributed to the slightly longer preparation protocol of the VAM printing process, which may have led to an initial delay in cell proliferation. By day 7, no significant difference was observed anymore between both methods (*p = 0.9054*). This suggests that the MSCs in the VAM-printed samples recovered and adapted to their environment, exhibiting a proliferation comparable with the film-cast samples. This is in line with other studies, where encapsulated cells needed a few days to adjust to the new environment.[73] At days 14 and 21, significant differences re-emerged (*p < 0.0001* for both time points). While for both groups, cells continued to proliferate, the film-cast samples exhibited a higher relative increase in cell number. This observation aligns with a growing body of literature highlighting the influence of hydrogel stiffness on cell behavior, particularly cell proliferation and - differentiation. For example, Tan *et al.* [74] demonstrated a correlation between hydrogel stiffness and cell fate, with softer materials tending to favor proliferation. Studies by Engler *et al.* [75] showed that matrix elasticity directed stem cell lineage specification, with softer substrates maintaining stemness and promoting proliferation. In our study, the film-cast samples, being less crosslinked and having a lower E-modulus than the VAM-printed ones (as discussed in section 3.3.2), provide a more compliant and potentially more porous environment facilitating cell expansion and nutrient and waste product exchange.

**Figure 7.**
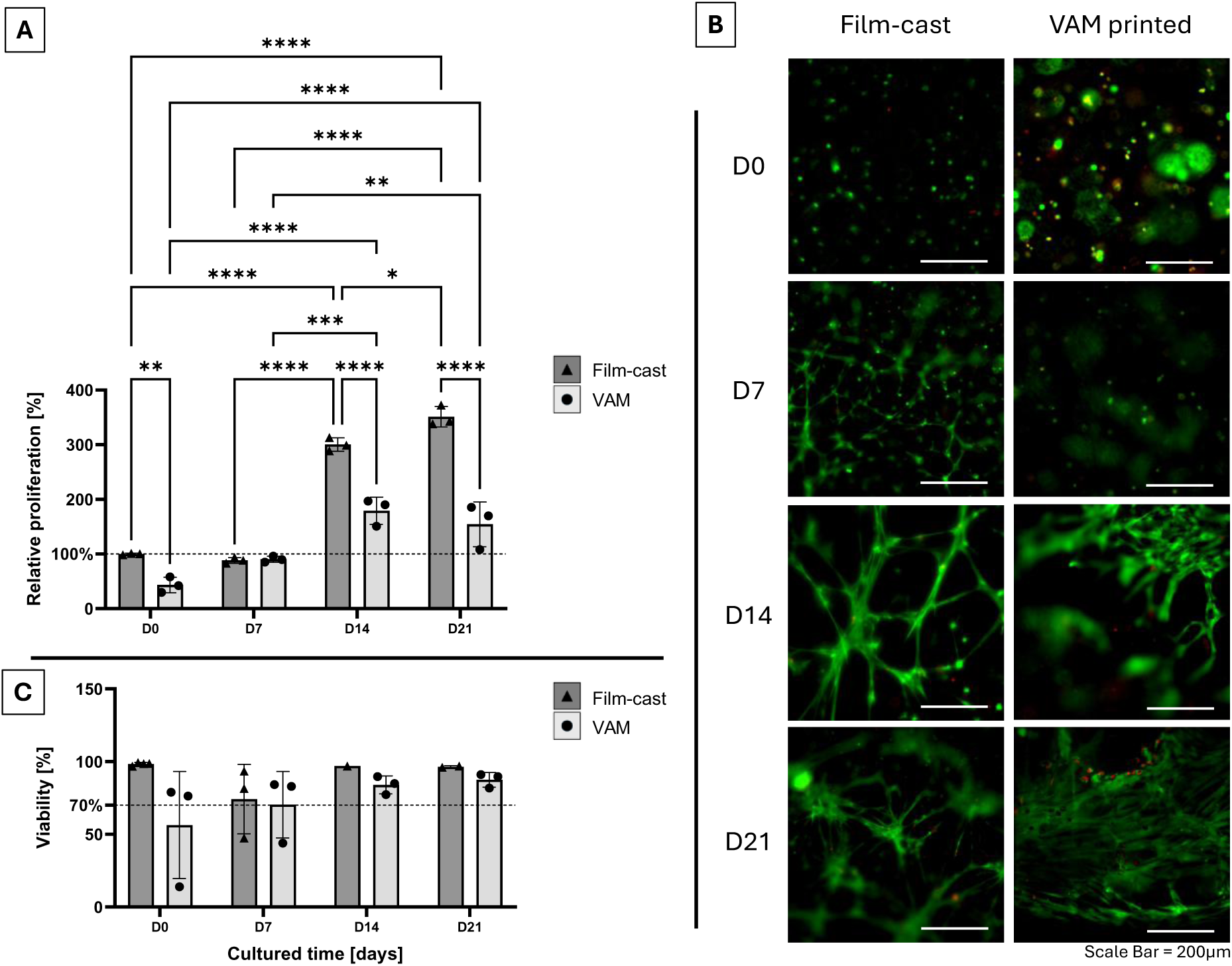
(A) Evaluation of proliferation (through an MTS assay) of MSC in VAM-printed versus film-cast encapsulation in GelNB-GelSH63 hydrogel (10% (w/v)) at different time points (day 0, 7, 14 and 21) (n = 3). The data was normalized against the D0 of the film-cast group (100%) and analyzed using a two-way ANOVA with Sidak’s multiple comparisons test. (* = p ≤ 0.05; ** = p ≤ 0.01; *** = p ≤ 0.001; **** = p ≤ 0.0001); (B) Viability (live/dead ratio) of both groups at the different time points; (C) Live/Dead (L/D) staining images of VAM-printed and film-cast MSCs at different time points (n = 3). Green indicates living cells, and red indicates dead cells. Scale bar: 200 μm.

However, it should be emphasized that the VAM-printed samples, despite their higher crosslinking density, still supported significant and sustained cell proliferation throughout the entire 21-day culture period, as evidenced by significant differences in absorbance between day 0 and later time points (D14 and D21, both *p < 0.0001*), and between day 7 and subsequent days (D14: *p = 0.0003*, D21: *p = 0.0059*). This is consistent with several studies demonstrating that cells can proliferate within even relatively stiff 3D-printed hydrogel constructs.[73,76] This finding highlights the versatility of VAM printing for creating scaffolds that can support cell viability and proliferation while also offering precise control over scaffold architecture. In the film-cast group, significant increases were similarly evident from day 0 to day 14 and day 21 (both *p < 0.0001*), and between day 7 and the later time points (both *p < 0.0001*).

Moreover, the increased stiffness of the VAM-printed constructs (see mechanical characterization in section 3.3.2), may offer advantages in terms of directing cell differentiation towards specific lineages, such as osteogenic pathways, as suggested in literature.[74,75,77] This potential for modulating cell fate through scaffold stiffness, underscores the importance of considering the interplay between material properties and cellular responses in designing scaffolds for tissue engineering applications.

The live/dead (L/D) staining images (Figure 7.B) provide further insights into cell morphology within the hydrogel constructs. Cells near the surface (in both VAM and film-cast samples) exhibited increased elongation over time, while those encapsulated deeper within the hydrogel maintained round. Despite these morphological differences, the MTS assay indicates that both populations remained viable and proliferative, suggesting that the microenvironment supports cell growth regardless of spatial positioning within the hydrogel. As a proof-of-concept to show the feasibility of printing porous scaffolds, the IWP design was volumetrically printed and the L/D was evaluated at day 3. Figure 8 shows the VAM-printed scaffold with mainly green-stained cells (living), as well as the difference in MSC morphology at the surface (elongated) and in the center (round-shaped) of the hydrogel. To validate the efficacy of the live/dead staining protocol, a representative control image showing both live and dead cells is included in the Supplementary Information (Figure S17).

**Figure 8.**
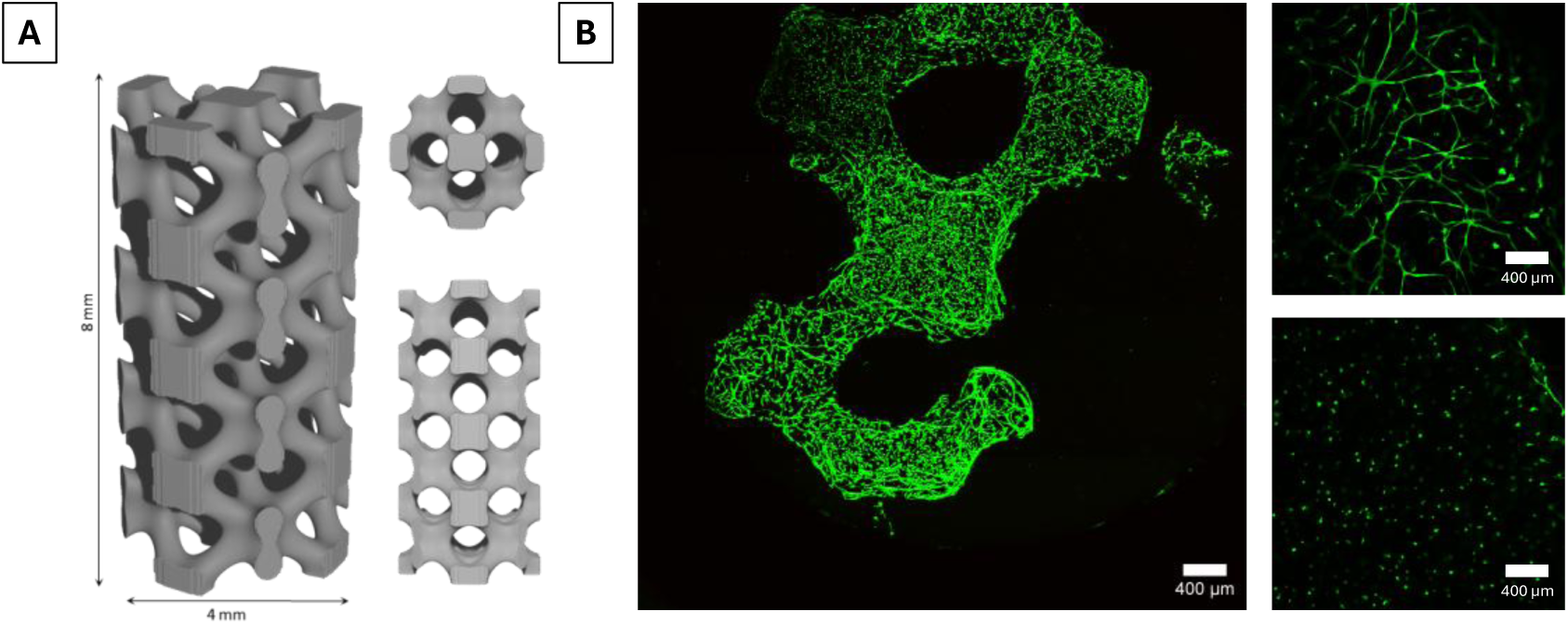
(A) CAD image of the IWP structure; (B) Live/Dead (L/D) staining images of VAM-printed MSCs at day 3. Green indicates living cells, and red indicates dead cells. Scale bar: 400 μm.

The structural integrity of the VAM-printed scaffolds was maintained during early cell culture, as shown in Figure 8 (Day 3), with no visual signs of deformation. Although long-term imaging has not been performed, VAM-printed discs cultured for 21 days under identical conditions also remained structurally intact upon visual inspection. This is consistent with findings by Parmentier *et al.* [4] who showed that gelatin-based VAM-printed scaffolds retained their architecture and supported cell growth over 21 days, confirming their stability for in vitro applications up to at least 21 days under the conditions used in this work.

#### 3.5.2 Differentiation assays

To confirm the identity of the MSCs and show the efficacy of the differentiation medium, MSCs were first differentiated towards osteogenic, chondrogenic and adipogenic lineage in 2D culture conditions [78–80] (Figure S18). For all assays film-cast samples cultured without differentiation factors (expansion medium) were used as 100% reference.

Osteogenic differentiation was evaluated using alkaline phosphatase activity (ALP) and calcium deposition, and compared between VAM-printed and film-cast hydrogels (Figure 9.A and 9.B). Additionally, to evaluate the need of providing extra growth factors to support differentiation, the cell response to differentiation (DIFF) and normal expansion (CTRL) medium was compared. Notably, the VAM-printed DIFF group displayed a significant increase in ALP activity at day 7 compared to their film- cast counterparts (vs CTRL: *p = 0.0350*, vs DIFF: *p = 0.0038*) indicating that VAM-printed scaffolds more effectively support early osteogenesis. This trend continued, with the VAM-printed scaffolds exhibiting markedly higher ALP activity after being cultured in DIFF medium for 21 days (*p < 0.0001*), suggesting an accelerated differentiation process within these constructs. An increase in ALP activity on stiffer gels has also been described by Mullen *et al.* [81]. Another study reported on significant changes in ALP activity with varying stiffness of the developed materials, with or without differentiation medium.[82]

**Figure 9.**
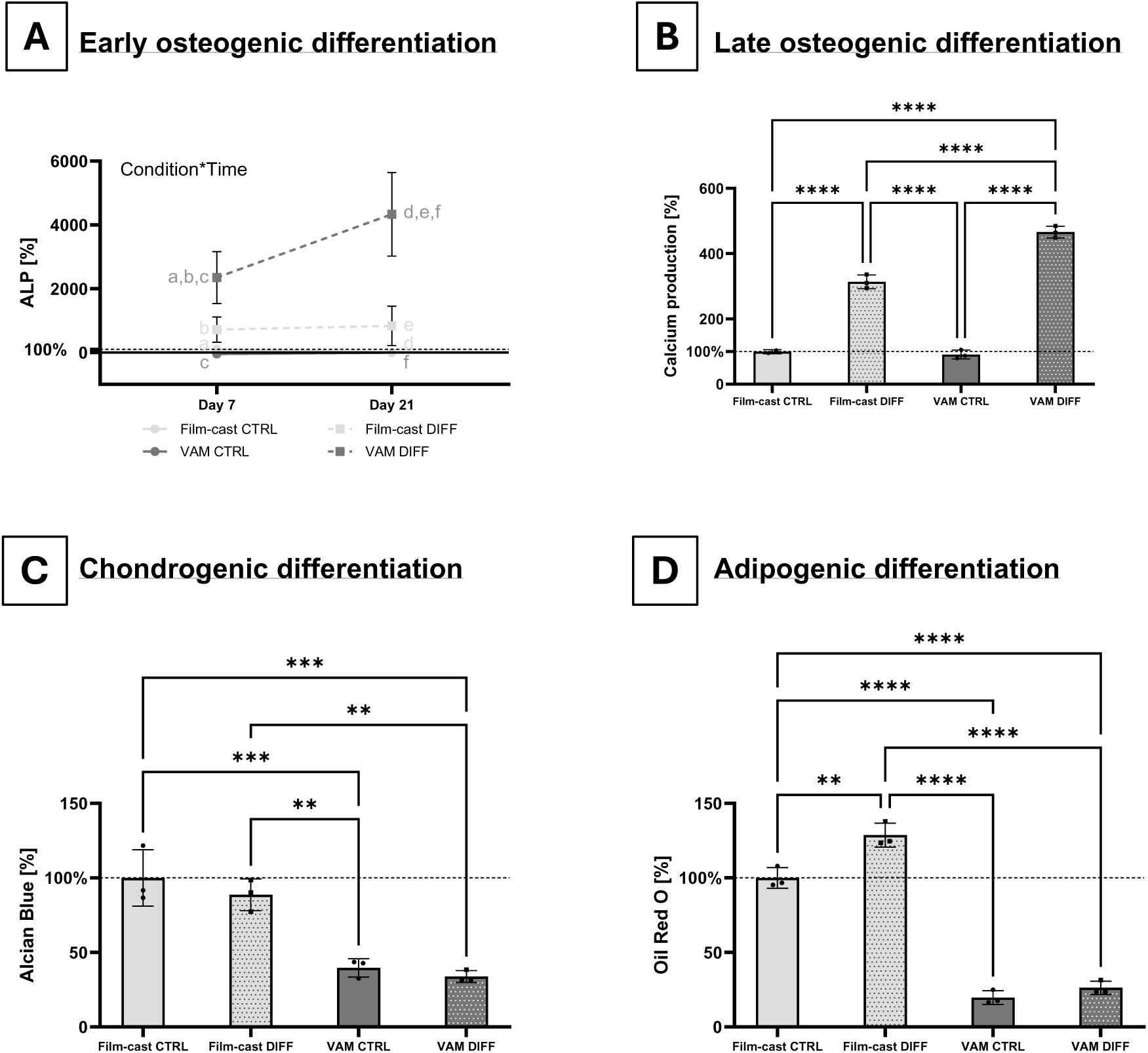
Results of osteogenic (ALP activity (A) and Ca deposition (B)), chondrogenic (Alcian Blue, (C)) and adipogenic (Oil Red O, (D)) differentiation of MSCs encapsulated in VAM-printed vs film-cast scaffolds (cultured in expansion (CTRL) or differentiation (DIFF) medium) (n = 3). The data was analyzed using a two-way ANOVA (ALP) or one-way ANOVA (Ca, Alcian Blue and Oil Red O) and a Tukey’s multiple comparisons test. In (A), the letters ‘a’ to ‘f’ indicate the significant differences of the two-way ANOVA test between the groups that share the same letter: a: **, b and c: *, d, e and f: ****. (* = p ≤ 0.05; ** = p ≤ 0.01; *** = p ≤ 0.001; **** = p ≤ 0.0001)

Analogously, calcium production on day 21 revealed significantly elevated levels in the differentiated VAM-printed group compared to the film-cast groups (*p < 0.0001*). The differentiated VAM-printed scaffolds showed the highest calcium deposition, indicative of enhanced late-stage osteogenesis. The sustained elevation in ALP activity alongside increased calcium levels in the VAM-printed scaffolds highlights the positive effect of VAM printing on osteogenesis. These findings suggest that the VAM- printed scaffolds create a more supportive environment for osteogenic differentiation, likely attributed to their more densely crosslinked network. It should also be noted that for both osteogenic differentiation assays, there is a significant increase in differentiation for the differentiated groups (DIFF) compared to the control groups (CTRL). This implies that osteogenic differentiation factors are required to induce differentiation, and that there is a synergistic effect between the differentiation medium and the processing method (VAM printing) in terms of osteogenic induction. The latter has also been described by Tan *et al.* [74] who showed that the interactions of chemical and physical factors may work synergistically to enhance bone regeneration. Other synergistic effects that have been described regarding osteogenic differentiation include among other surface chemistry and surface topography [83], cell-derived extracellular matrices and topography [84], stiffness and structure of 3D-printed hydrogels [77], hydrogel stiffness and co-culture systems [54], surface topography and induction media [85].

Chondrogenic differentiation was evaluated using the Alcian Blue staining to assess the presence of GAGs (Figure 9.C).[86–88] When comparing chondrogenic potential in the differentiated versus control samples, no significant difference in GAG production could be observed – though positive responses indicated the presence of GAGs in both groups. This can be explained by the fact that the hydrogel scaffold might induce the chondrogenic differentiation of MSCs, irrespective of the used medium. It must also be mentioned that spontaneous chondrogenic differentiation of equine MSCs has been reported before [78,89], and was hypothesized to be linked to high-density culture conditions and hypoxia in the 3D cell culture system. This spontaneous chondrogenic differentiation potential has also been observed in human [90,91] and bovine MSCs [92,93]. Nevertheless, this assay further emphasizes the differences in GAG production between VAM-printed and film-cast scaffolds. The GAG production in the VAM-printed groups significantly decreased (*p < 0.05*) compared to the film-cast group. In a study by Goldshmid *et al.* [94] it was shown that chondrogenesis was favored in materials with a higher modulus (i.e. G’ = 1000 Pa). It should be noted that in this study, different concentrations of PEG-diacrylate were used to modify the stiffness of a bovine PEG-fibrinogen hydrogel, meaning both mechanical and chemical properties could have influenced the observed chondrogenic response. Similarly, Zhou *et al.* [95] investigated the chondrogenic differentiation of MSCs using polyacrylamide-based hydrogels of varying stiffness (ranging from ∼0.5 kPa to stiffer formulations), prepared by altering the acrylamide-to-bis-acrylamide ratio. They showed that the chondrogenic differentiation of MSCs is promoted by soft substrates (about 0.5 kPa). Since these studies compared different hydrogel compositions rather than a single composition with varying mechanical properties (as in our study), the observed effects on chondrogenesis could possibly be attributed to mechanical as well as chemical differences. In addition, further analysis indicated that TGF-β3 supplementation increased the expression level of cartilage-related markers and masked the stiffness-derived expression pattern of hypertrophic markers.[95] Adipogenic differentiation of MSCs was assessed using the Oil Red O staining, which quantifies the lipid droplets intracellularly (Figure 9.D).[96–98] A robust lipid accumulation was observed in the film- cast scaffolds cultured in differentiation medium (DIFF), indicating successful adipogenesis. The results revealed again distinct outcomes between VAM-printed and film-cast scaffolds. In general, less adipogenic differentiation was observed in the VAM-printed group (*p < 0.0001*) as compared to the film-cast group, regardless of the media used. In a study by Zhao *et al.* [99], where hydrogels with varying stiffness have been evaluated (from 0.15 to 4 kPa) using Oil Red O staining, the MSCs exhibited a smaller spreading area with much more stretched morphology on the 4 kPa hydrogels than on the hydrogels with lower stiffness. The finding that substrate elasticity plays a crucial role in directing stem cell differentiation towards specific lineages also applies to adipogenic differentiation.[100] For instance, soft substrates (E = 2 kPa) promote adipogenic differentiation [34], whereas stiff substrates (E = 34 kPa) favor osteogenic differentiation [75]. Wen *et al.* [101] identified substrate elasticity as the primary factor influencing cell differentiation under unstrained culture conditions. In our study, the VAM-printed scaffolds showed an E of 21.04 ± 2.14 kPa (section 3.3.2) which is a 10-fold higher than the soft substrates of 2 kPa that promote adipogenic differentiation as described by Murphy *et al.* [34], which clearly explains the lower adipogenic differentiation results observed in the VAM-printed scaffolds.

The link between the mechanical properties of the VAM-printed scaffolds and their ability to support cell differentiation is pivotal in understanding these results.[102] As described in the section on the mechanical characterization of the VAM-printed scaffolds (section 3.3.2), a more efficient and denser crosslinking led to an E modulus of 21.04 ± 2.14 kPa that was 3-4 times higher than the film-cast (E = 6.467 ± 0.154 kPa). This resulted in the increased osteogenesis observed in the VAM-printed scaffolds, while the lesser crosslinked film-cast constructs supported the adipogenic and chondrogenic potential of encapsulated MSCs. It should also be mentioned that it is never one factor alone that is influencing MSC differentiation, but a combination of factors, e.g. substrate stiffness and surface topography [103], physico-mechanical and biomolecular cues [104], effect of matrix stiffness combined with TGF- β [100] or protein tethering [101].

In summary, we illustrated distinct differentiation outcomes for MSCs cultured in VAM-printed versus film-cast constructs. The VAM-printed scaffolds exhibited significantly higher ALP activity and calcium deposition, confirming effective osteogenesis. Conversely, chondrogenic and adipogenic differentiation, assessed through Alcian Blue and Oil Red O staining, were more pronounced in the film-cast groups. The inherent differences in crosslinking density and mechanical properties between VAM-printed and film-cast scaffolds are significant. While not demonstrated in this study, it is possible to obtain softer gels through VAM by modifying the chemical composition, such as the degree of substitution of the modified gelatins or the gelatin concentration of the bioresin. Additionally, film casting with alternative protocols, such as extended irradiation or post-curing, could be used to yield stiffer constructs, potentially targeting osteogenic applications. However, when comparing both techniques regarding their potential to fabricate 3D scaffolds, only VAM printing enabled the creation of complex structures, offering greater design flexibility. This study lays the foundation for future research on optimizing material formulation within the VAM printing framework to enhance differentiation across multiple lineages, related to the specific tissue engineering applications.

## 4. Conclusions

This study highlights the potential of photo-crosslinkable gelatin hydrogels (GelSH and GelNB) for volumetric additive manufacturing, with a focus on their ability to guide MSC fate and differentiation for tissue engineering applications. Material characterization revealed a clear relationship between the degree of substitution and the mechanical properties: GelSH functionalized with 1, 3, and 5 equivalents exhibited DS values of 39%, 54%, and 63%, respectively, while increasing the hydrogel concentration from 5 to 10% (w/v) led to a corresponding increase in the storage modulus (G’). This tunable behavior, with G’ ranging from 206 Pa to 12.5 kPa, highlights the versatility of these hydrogels in replicating diverse tissue environments.

When applied in VAM-printed constructs using GelNB-GelSH63 at 10% (w/v), the scaffolds demonstrated enhanced mechanical performance, with compressive strength reaching 507.63 ± 53.15 kPa and a compressive modulus of 21.04 ± 2.14 kPa. This structural integrity translated into significant biological benefits. VAM-printed scaffolds supported significant and sustained cell proliferation throughout the entire 21-day culture period. The more densely crosslinked network provided by the VAM-printed hydrogels also promoted osteogenic differentiation, as indicated by elevated ALP activity and calcium deposition when differentiation medium is used. In contrast, film-cast hydrogels, with their softer and less dense crosslinked structure, favored chondrogenic and adipogenic differentiation. These findings underscore the critical role of scaffold physico-chemical properties in guiding cell behavior, demonstrating the potential of VAM to create customized hydrogel scaffolds for targeted tissue engineering applications.

## Acknowledgement and funding

1. N. Pien, B. Bogaert and M. Meeremans would like to acknowledge the financial support of the Research Foundation Flanders (FWO) under the form of an FWO junior post-doctoral research grant (12E4523N), and FWO SB PhD fellowships (1SHE724N and 1S02822N), respectively. S. Van Vlierberghe acknowledges Interreg Healthy Teeth and the Research Foundation Flanders for providing a Hercules grant (I003922N).

## Notes

### Competing Interest Statement

The authors have declared no competing interest.

### Summary of Updates

This version has been revised to update text and figures according to the reviewer's comments and feedback.

