## Supplementary Information for "Exploring the Impact of Volumetric Additive Manufacturing of Photo-crosslinkable Gelatin on Mesenchymal Stromal Cell Behavior and Differentiation"

**Material synthesis**

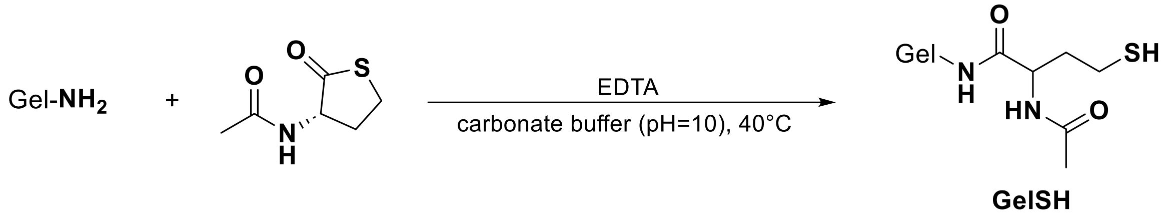


*Figure S1. Reaction scheme of thiolated gelatin.*

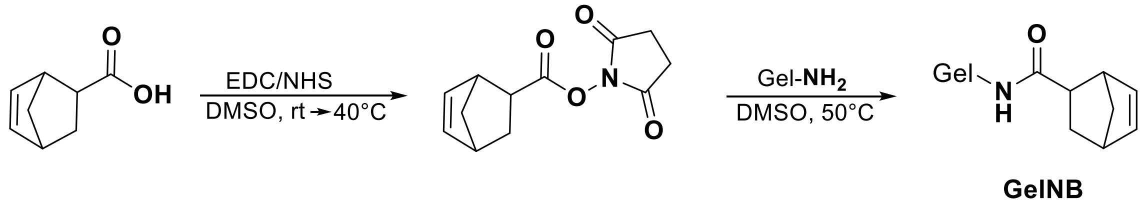


*Figure S2. Reaction scheme of norbornene-modified gelatin.*

*Table S1. Overview of the quantities used for synthesis of the thiolated gelatin with varying degree of substitution.*

| | *Gelatin [g]* | *Solvent [mL]* | *N-acetyl homocysteine thiolactone [g]* |
|---|---|---|---|
| *GelSH1eq* | 20 | 200 (carbonate buffer) | 1.225 |
| *GelSH3eq* | 20 | 200 (carbonate buffer) | 3.671 |
| *GelSH5eq* | 20 | 200 (carbonate buffer) | 6.193 |

The carbonate buffer is composed of 0.6721 g of $NaHCO_3$ and 1.2719 g of $Na_2CO_3$ dissolved in 200 mL of Milli-Q and pH adjusted to a pH = 10 ± 0.1 to afford a 0.1M buffer.

### OPA analysis

The borate buffer is composed of 0.6183 g of boric acid and 0.7455 g of potassium chloride dissolved in 100 mL of Milli-Q and pH adjusted to pH = 10 ± 0.1 to afford a 0.1M buffer.

| | *Blanc* | *0.002 mM* | *0.006 mM* | *0.01 mM* | *Gelatin B* | *GelSH1eq* | *GelSH3eq* | *GelSH5eq* | *GelNB* |
|---|---|---|---|---|---|---|---|---|---|
| *M1* | 0.2293 | 0.3589 | 0.6511 | 0.9267 | 0.8149 | 0.5549 | 0.5009 | 0.4448 | 0.4103 |
| *M2* | 0.2652 | 0.3993 | 0.5899 | 0.6862 | 0.8185 | 0.6040 | 0.4972 | 0.4386 | 0.4543 |
| *M3* | 0.2268 | 0.3398 | 0.5493 | 0.8815 | 0.7503 | 0.5819 | 0.4972 | 0.4631 | 0.4115 |

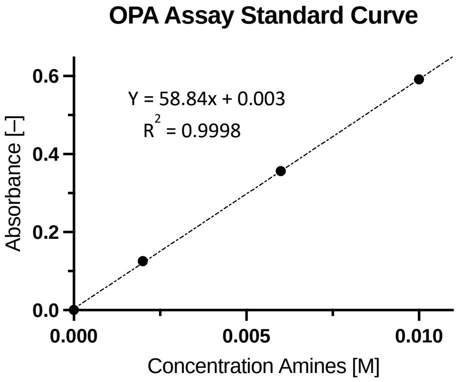


| | *GelSH1eq* | *GelSH3eq* | *GelSH5eq* | *GelNB* |
|---|---|---|---|---|
| *DS1 [%]* | 43.5256 | 53.3325 | 63.5208 | 69.7863 |
| *DS2 [%]* | 34.6086 | 54.0045 | 64.6468 | 61.7955 |
| *DS3 [%]* | 38.6222 | 64.0045 | 60.1973 | 69.5684 |
| *Average DS [%]* | 38.9188 | 53.7805 | 62.7883 | 67.0501 |
| *STDEV DS [%]* | 4.4659 | 0.3879 | 2.3134 | 4.5519 |

$$n_{NH_2}[mmol \cdot g^{-1}] = \frac{\frac{(ABS - BLANK) - b}{a}}{C}$$

With "*ABS*" being the absorbance values measured with UV-VIS, "*BLANK*" the average of the blank measurements, "*a*" and "*b*" obtained from the calibration curve (y= ax+b) as indicated on the Figure, and "*C*" being the concentration in $mg \cdot mL^{-1}$.

$$\text{DS [\%]} = \frac{n_{NH_2}\, gelatin - n_{NH_2}\, modified\ gelatin}{n_{NH_2}\, modified\ gelatin} \cdot 100$$

$n_{NH2}$ gelatin = amount of free amines in (unmodified) gelatin, expressed in $mmol \cdot g^{-1}$

$n_{NH2}$ modified gelatin = amount of free amines in modified gelatin, expressed in $mmol \cdot g^{-1}$

**LAP synthesis**

Lithium phenyl-2,4,6-trimethylbenzoylphosphinate (LAP) was synthesized in-house by dissolving 9.45 g (109 mmol, 4 equivalents) in 150 mL of butan-2-one at 65°C. Next, 8.6 g of TPO-L (27.2 mmol, 1 equivalent) was added to the mixture and allowed to stir for 24h at 65°C. During these 24h a white precipitate formed which was filtered by vacuum filtration and rinsed with 2x 150mL of petroleum ether.

*Table S2. Hydrogel formulations for preparation of (a) hydrogel films and (b) VAM photo-resins (without MSCs) and bioresins (with 2·10⁶ MSCs per mL).*

| *(a)* | *GelNB* | *GelSH* | *Concentrations* | *PI* | *Solvent* |
|---|---|---|---|---|---|
| *GelNB-GelSH**39*** | 1.2 eq = DS 60% | 1 eq = **DS 39%** | 5, 7.5 and 10% (w/v) in equimolar ratio NB to SH | 0.0076% (w/v) LAP for films | Ultrapure water |
| *GelNB-GelSH**54*** | 1.2 eq = DS 60% | 3 eq = **DS 54%** | 5, 7.5 and 10% (w/v) in equimolar ratio NB to SH | 0.0076% (w/v) LAP for films | Ultrapure water |
| *GelNB-GelSH**63*** | 1.2 eq = DS 60% | 5eq = **DS 63%** | 5, 7.5 and 10% (w/v) in equimolar ratio NB to SH | 0.0076% (w/v) LAP for films | Ultrapure water |

| *(b)* | *GelNB* | *GelSH* | *Concentrations* | *PI* | *Solvent* | *Cells* | *Iodixanol* |
|---|---|---|---|---|---|---|---|
| *GelNB-GelSH**63*** | 1.2 eq = DS 60% | 5eq = **DS 63%** | 10% (w/v) GelNB + GelSH in equimolar ratio NB to SH | 0.05% (w/v) LAP for VAM | DPBS | - | - |
| *GelNB-GelSH**63** (with MSC)* | 1.2 eq = DS 60% | 5eq = **DS 63%** | 10% (w/v) GelNB + GelSH in equimolar ratio NB to SH | 0.05% (w/v) LAP for VAM | DPBS | $2 \cdot 10^6$ MSCs per mL | 0, 10, 15, 20% (v/v) |
| *GelNB-GelSH**63** (optimized bioresin)* | 1.2 eq = DS 60% | 5eq = **DS 63%** | 10% (w/v) GelNB + GelSH in equimolar ratio NB to SH | 0.05% (w/v) LAP for VAM | DPBS | $2 \cdot 10^6$ MSCs per mL | 15% (v/v) |

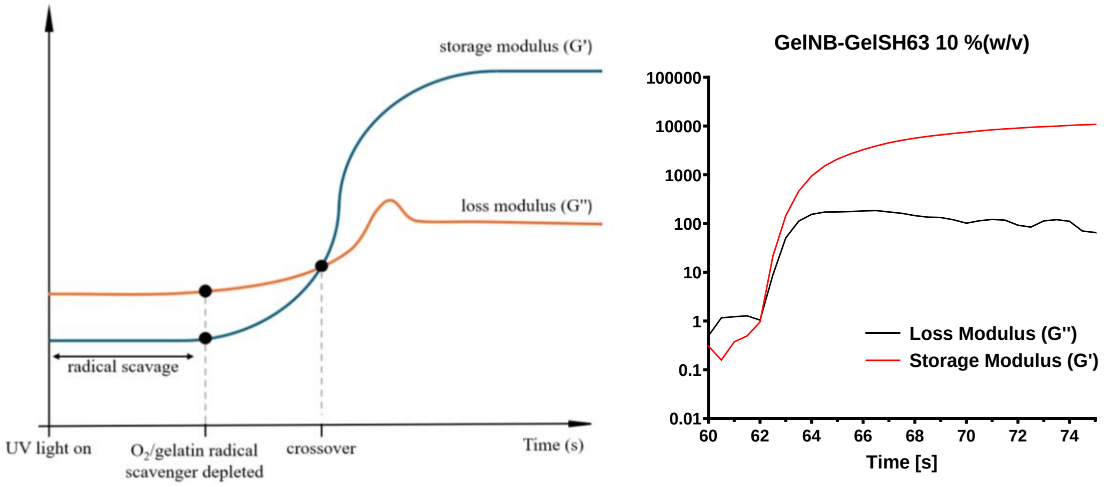


*Figure S3. Illustration of crossover point identification in photo-rheology*

### S1. Development of photo-crosslinkable gelatins and determination of degree of substitution

The development of thiolated gelatin (GelSH) and gelatin-norbornene (GelNB) yielded biomaterials, each with functional groups optimized for photo-crosslinking.[1] GelSH was obtained through modification of gelatin with N-acetyl-homocysteine thiolactone to introduce thiol groups, while GelNB was obtained through gelatin functionalization with norbornene groups to enable rapid photo-crosslinking via thiol-ene chemistry under light exposure.[2,3] These modifications were carefully controlled to achieve specific degrees of substitution that allow tunable mechanical properties. The selection of GelSH and GelNB was motivated by their ability to undergo fast, efficient polymerization [4], which is critical for the precision and structural integrity required in volumetric additive manufacturing (VAM), especially for creating complex 3D hydrogel structures suitable for tissue engineering applications.[5–7]

An OPA assay was employed to assess the functionalization levels of thiolated gelatin as a function of the equivalents of N-acetyl homocysteine thiolactone (1eq, 3eq, and 5eq) and of norbornene-modified gelatin. The results indicated that the DS of GelSH5eq was the highest with 62.8 ± 2.3% followed by GelSH3eq being 53.8 ± 0.4% and lastly GelSH1eq with a DS of 38.9 ± 4.5%. For GelNB, only one variant was developed (with a targeted DS of 60%). The DS was determined both by the OPA assay and $^{1}$H-NMR spectroscopy (Figure S4). The OPA assay yielded a DS of 67.1 ± 4.6% whereas the latter yielded a lower DS of 55%. These results are in line with previously reported data on GelSH and GelNB (DS of 67-72% for GelSH5eq and DS of 53% via NMR spectroscopy for GelNB1.2eq) [2,8] and it can thus be concluded that the materials were successfully functionalized. Due to the overlap of the NMR signals of norbornene and N-acetyl-L-cysteine with the reference signal of gelatin (at 1 ppm), it is difficult to accurately quantify the DS using $^1$H-NMR [9], which can also explain the differences in DS when comparing the results of $^1$H-NMR spectroscopy and the OPA assay. For further annotations in the manuscript, we refer to the developed materials (based on the DS determined via OPA) as GELSH39, GELSH54 and GELSH63, for the 1, 3 and 5 eq, respectively. Because only 1 variant of GELNB was developed (i.e. 1.2 eq), we will simply refer to it as GELNB.

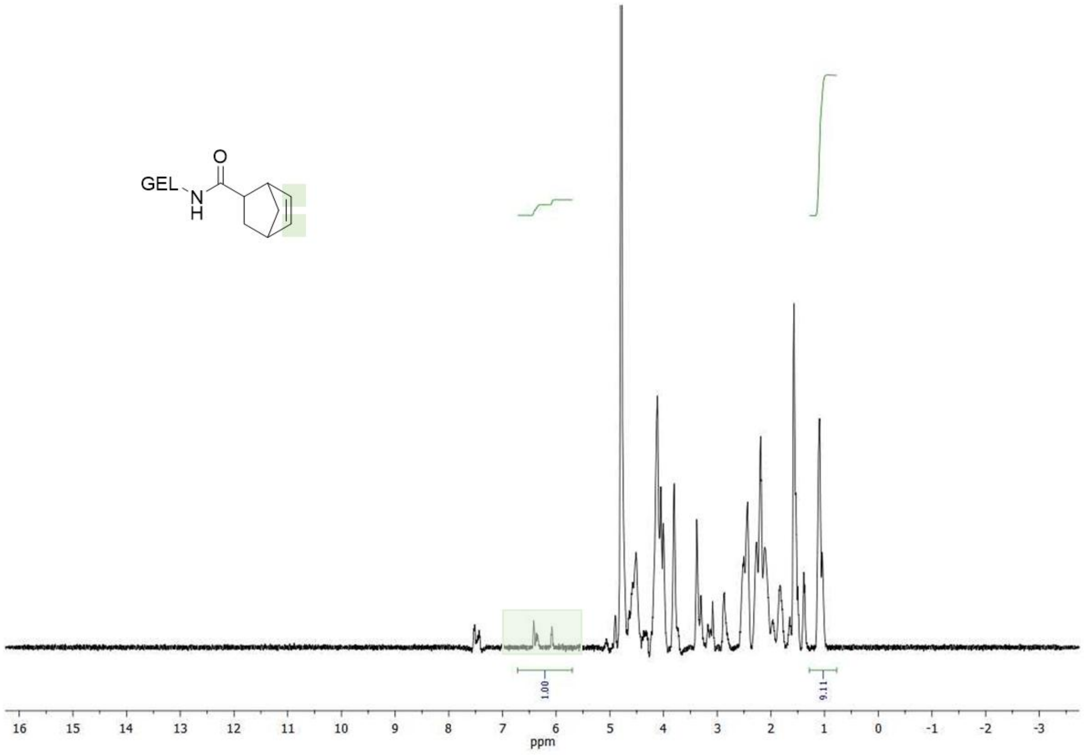


*Figure S4. $^1$H-NMR spectrum of GelNB1.2eq*

### S2. In situ hydrogel crosslinking behavior via photorheology

Photorheology was performed to investigate and compare the cross-linking behavior and the mechanical properties of the GelNB-GelSH hydrogels at varying concentrations (5, 7.5, and 10% (w/v)) and modification degrees (39%, 54% and 63%). The storage modulus (G'), which indicates the mechanical strength of the gels was evaluated. The values of G′ can vary significantly based on the specific formulation and crosslinking conditions employed. In studies involving gelatin-based hydrogels, it has been observed that the storage modulus is influenced by the degree of substitution and the concentration of the gelatin precursors applied. For instance, it has been reported that increasing the degree of substitution in gelatin hydrogels leads to a higher storage modulus due to a denser and stiffer network structure.[2] Similarly, the mechanical properties of GelNB hydrogels can be tuned by adjusting the concentration of the thiol and norbornene components, which directly affects the crosslinking density and consequently the G′ values.[10] Moreover, the rheological properties of GelNB-GelSH hydrogels have already been characterized in various studies. For example, the storage modulus of GelNB-based hydrogels has been reported to range from approximately 0.2 to 17 kPa depending on the crosslinking density and formulation.[11–14] This variability underscores the tunability of the mechanical properties of these hydrogels, which is essential for applications in tissue engineering and regenerative medicine.

Our results (Figure S5) clearly showed a trend where increasing the concentration of GelNB-GelSH resulted in a higher storage modulus. For example, GelNB-GelSH63 at 10% (w/v) exhibited the highest G' values (~12.5 kPa), significantly outperforming the GelNB-GelSH63 formulations at lower concentration, i.e. 5 and 7.5% (w/v). Similarly, a lower modification degree (e.g., GelNB-GelSH39) resulted in weaker gels, as demonstrated by consistently lower G' values compared to the GelNB-GelSH63 formulations. The significant differences in mechanical properties between the different formulations ($p < 0.01$) stress the importance of optimizing both the concentration and modification degree of the hydrogels for specific tissue engineering applications. Stiffer hydrogels, characterized by higher storage moduli (G'), have been widely reported to promote osteogenic differentiation, particularly by providing mechanical cues that resemble the rigidity of bone tissue. While these hydrogels do not fully replicate the mechanical properties of native bone - which typically exhibits moduli in the GPa range - they offer a stiffer microenvironment relative to softer hydrogels, which can influence stem cell fate. More specifically, bone TE studies have reported G' values in the kPa to MPa range, depending on the hydrogel composition and cell type.[13,15,16] Hydrogels with intermediate G' values, typically in the kPa range, are often more conducive to chondrogenesis.[17–19] This reflects the softer and more flexible nature of cartilage compared to bone. Softer hydrogels with lower G' values, typically in the < 2 kPa range, tend to favor adipogenesis.[2,20,21] This mimics the compliant environment of adipose tissue. Comparing these literature data to our findings (Figure 3), our GelNB-GelSH63 formulation at 10% (w/v) exhibited a G' of ~12.5 kPa, which is situated within the lower end of the range for osteogenesis, while the G' values of our other formulations, particularly at higher concentrations, align with the kPa range associated with chondrogenesis and adipogenesis.

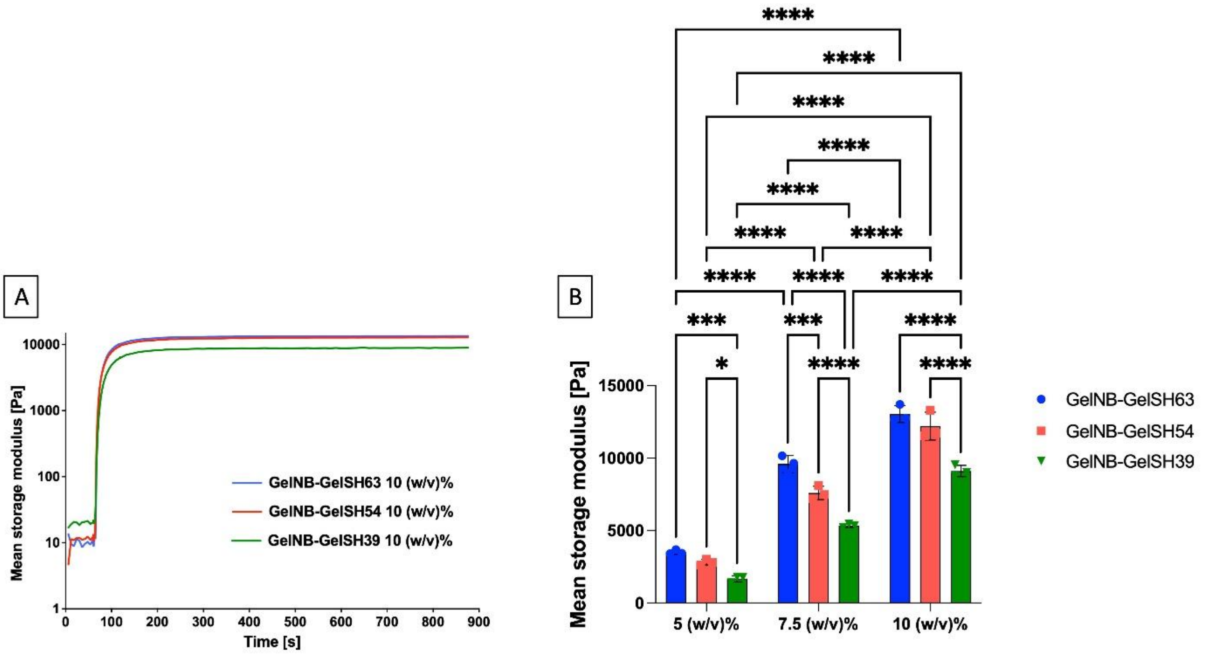


*Figure S5. (A) Photorheology comparison of GelNB-GelSH39, 54 and 63 (i.e. 1eq, 3eq and 5eq) for 10% (w/v), all with 0.0076% (w/v) photo-initiator. (B) Plateau values of storage moduli (G'), measured by photorheology, of GelNB-GelSH solutions (1 eq, 3eq and 5eq for 5, 7.5 and 10% (w/v)). Data (n = 3) was analyzed using a two-way ANOVA and a Tukey's multiple comparisons test. (* = p ≤ 0.05; ** = p ≤ 0.01; *** = p ≤ 0.001; **** = p ≤ 0.0001)*

### S3. Determination of hydrogel gel fraction and mass swelling ratio

In terms of gel fraction, gelatin-based hydrogels generally exhibit high gel fractions, often reported to be around 90% or higher.[2,13] Across all concentrations and combinations used in this work, the gel fraction is close to 100% (Figure S6). This indicates both proper crosslinking, as very few unmodified or uncrosslinked chains remain in the hydrogel as well as little to no leaching of potentially harmful compounds. With regard to cell viability, another important characteristic is the hydrogel's capacity to mimic the highly aqueous extra-cellular matrix (ECM) of tissues. To this end, the mass swelling ratio q was determined by evaluating the difference between the dried state and the swollen state of the gels. The mass swelling ratios ranged between 20 and 60 for 10% (w/v) to 5% (w/v) concentrations, being consistent with previously reported values for similar hydrogel systems.[2,3,8,13,14] The swelling behavior of GelNB-GelSH hydrogels is attributed to the hydrophilic nature of gelatin and the crosslinked network structure, which allows for significant water uptake while maintaining structural integrity.[22] While these swelling ratios indicate the formation of robust hydrogels, their suitability for optimal cell response depends on the specific cell type and anticipated application.[23,24] The swelling ratio can be influenced by various factors, including the degree of crosslinking and the specific formulation of the hydrogel. For instance, hydrogels with higher crosslinking densities tend to exhibit lower swelling ratios due to the reduced mobility of polymer chains, which limits the amount of water that can be absorbed.[2,25] Generally, it can be observed that the mass swelling ratio decreases with an increasing concentration of the material. This highlights the complex interplay between DS, concentration, and network properties in GelNB-GelSH hydrogels.

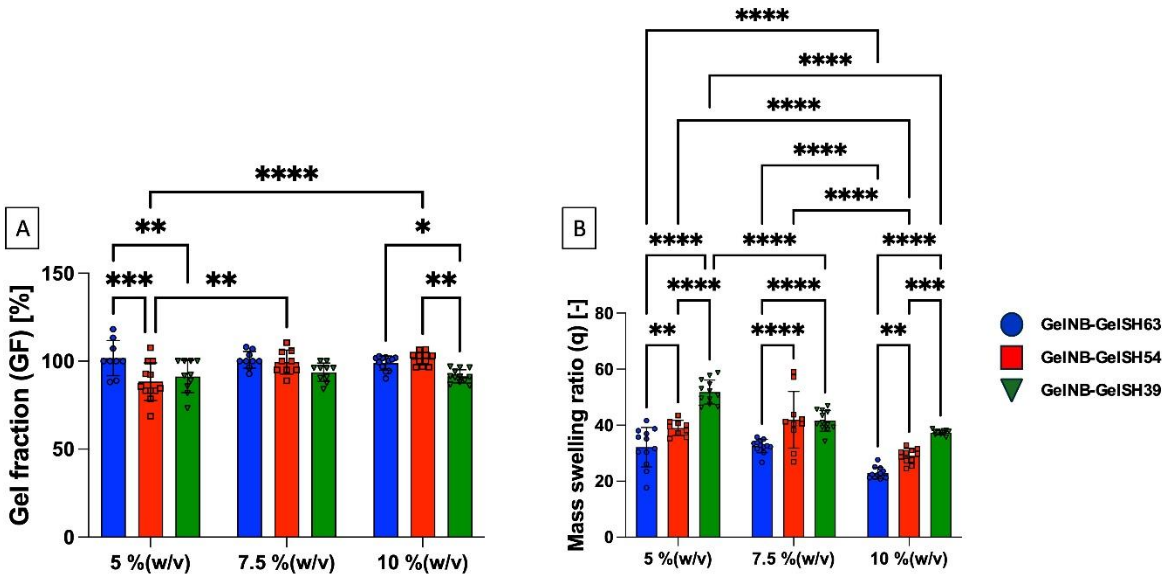


*Figure S6. Gel fraction (A) and mass swelling ratio (B) on the various gelatin formulations (GelSH with a DS of 63, 54 and 39, GelNB-GelSH concentrations of 10, 7.5 and 5% (w/v), n = 6). A two-way ANOVA and a Tukey's multiple comparisons test was selected. (* = p ≤ 0.05; ** = p ≤ 0.01; *** = p ≤ 0.001; **** = p ≤ 0.0001)*

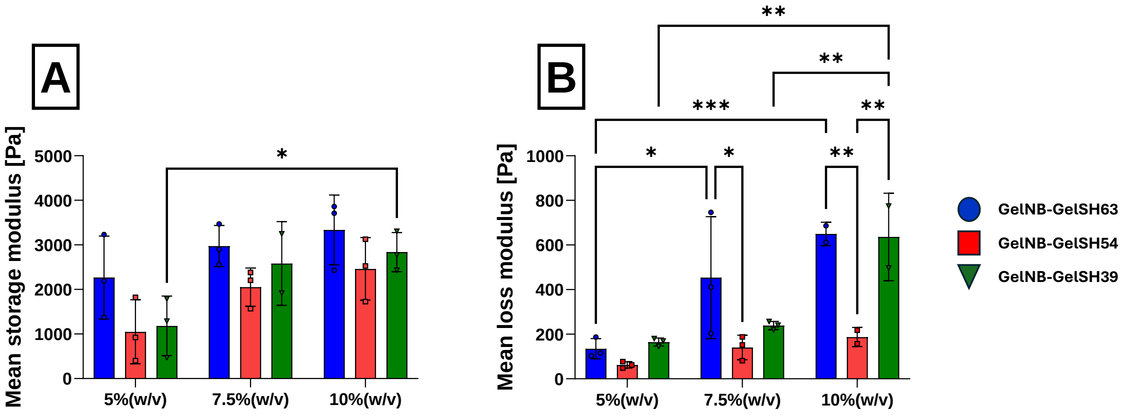


*Figure S7. Frequency sweep analysis on crosslinked GelNB-GelSH (after equilibrium swelling) (n = 3). A two-way ANOVA and a Tukey's multiple comparisons test was selected. (* = p ≤ 0.05; ** = p ≤ 0.01; *** = p ≤ 0.001; **** = p ≤ 0.0001)*

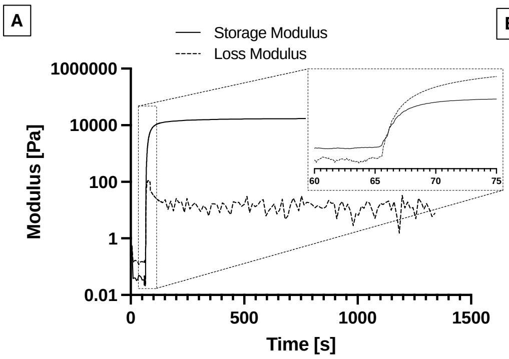


B

| Intensity [mW·cm⁻²] \ Time [s] | 5.83 | 7.12 | 8.69 | 10.62 | 12.96 |
|---|---|---|---|---|---|
| 6.48 | Small Dot | Large Dot | Large Dot | Large Dot | Large Dot |
| 5.31 | Small Dot | Large Dot | Large Dot | Large Dot | Large Dot |
| 4.35 | Barely Visible Dot | Large Dot | Large Dot | Large Dot | Large Dot |
| 3.56 | No Dot | Barely Visible Dot | Small Dot | Large Dot | Large Dot |
| 2.92 | No Dot | No Dot | Barely Visible Dot | Barely Visible Dot | Barely Visible Dot |

*Figure S8. Determination of the reference dose for VAM printing. (A) Photo-rheology analysis to determine the gel point and required light dose for hydrogel crosslinking. The storage modulus (solid line) and loss modulus (dashed line) were monitored over time, with a gelation point observed at the crossover. The inset highlights the modulus increase during the gelation process. (B) Evaluation of reference doses using the Readily3D protocol, showing the relationship between light intensity (mW·cm$^{-2}$) and exposure time (s) to achieve effective crosslinking.*

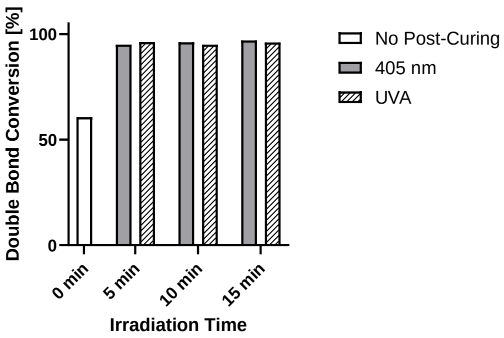


*Figure S9. Post processing curing of GelNB-GelSH63 10% (w/v) after VAM printing influences double bond conversion (measured by HR-MAS).*

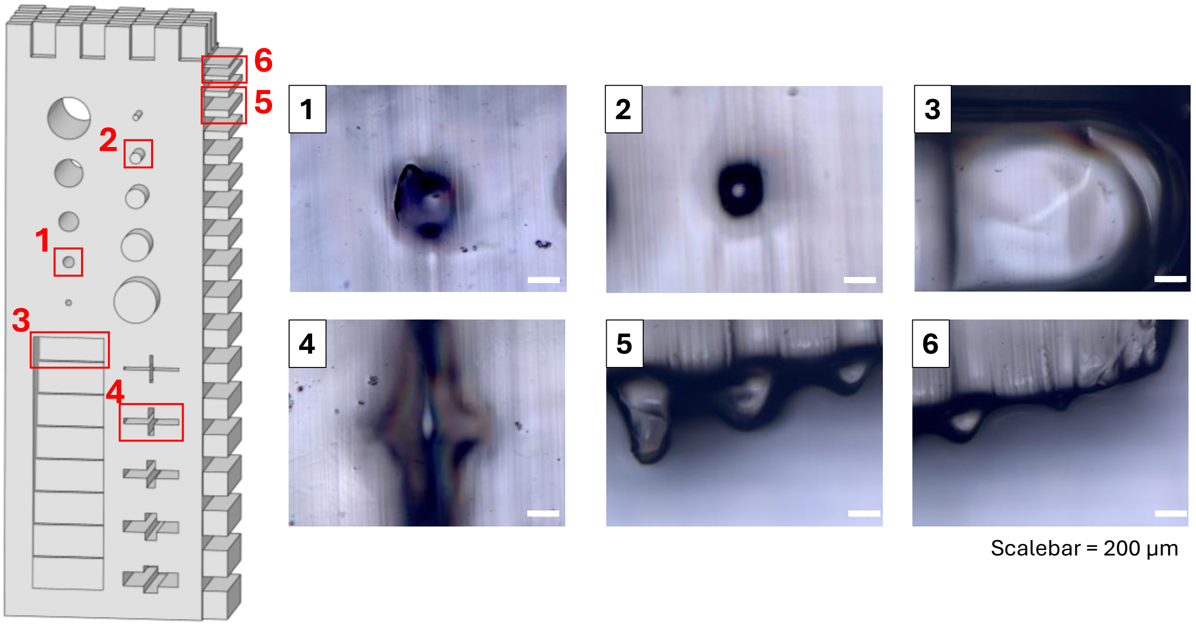


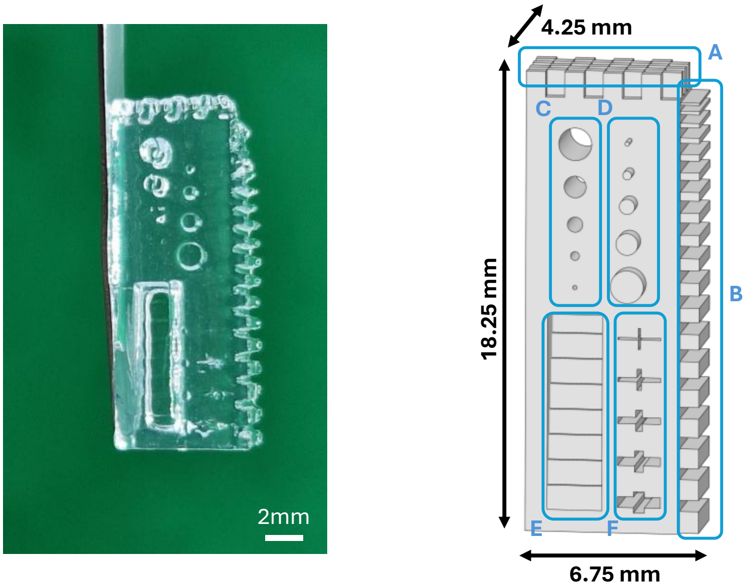


- **Feature A:** Cuboid checkered pattern with 0.75x0.5x1 mm
- **Feature B:** Walls with length height of 0.75mm ranging from 0.085 to 0.85 mm
- **Feature C:** Protruding holes with diameter of 0.17, 0.34, 0.60, 0.86 and 1.29 mm
- **Feature D:** Cylinders with diameter of 0.17, 0.34, 0.60, 0.86 and 1.29 mm and height of 0.83
- **Feature E:** Stair steps than thin out from 1.72 to 0.21 mm
- **Feature F:** Protruding crosses ranging thickness from 0.08 to 0.43 mm

*Figure S10. Top: CAD/CAM mimicry measured using optical light microscopy immediately after VAM printing. Bottom: Photograph showing the VAM printed structure.*

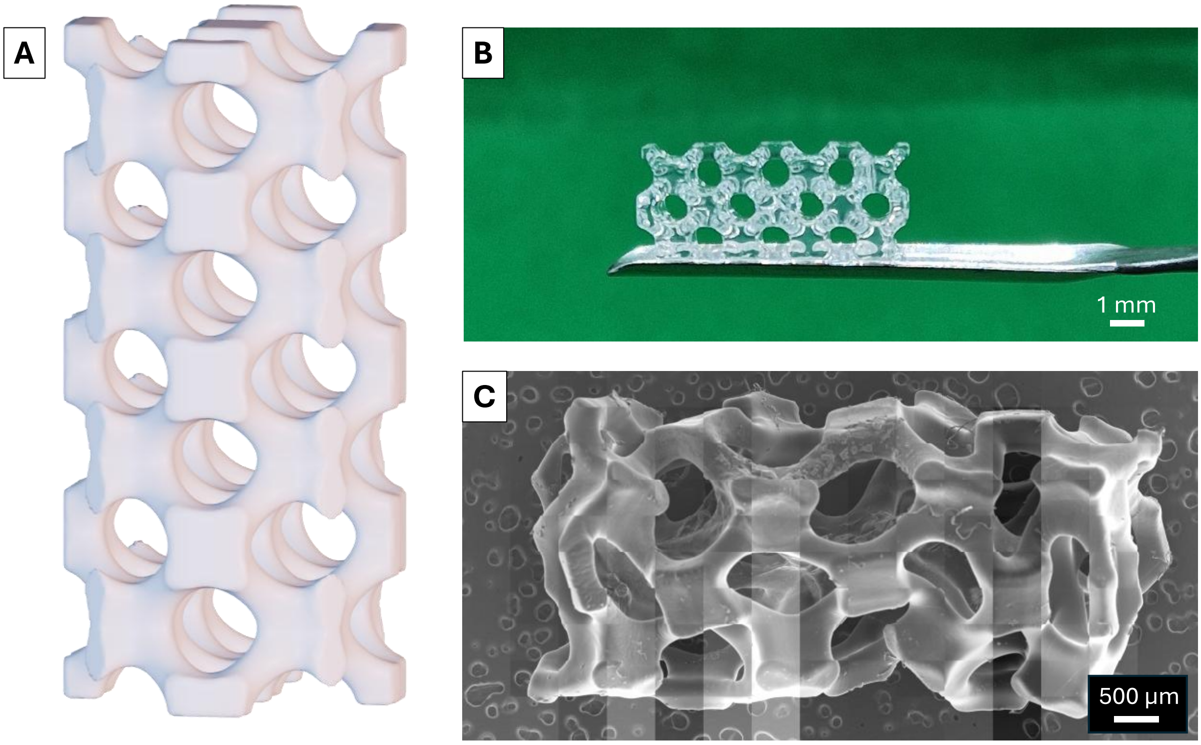


*Figure S11. CAD/CAM mimicry of stent-like structure, VAM printed in GelNB-GelSH63 10% (w/v) (0.05% (w/v) LAP), obtained via optical light microscopy immediately after printing.*

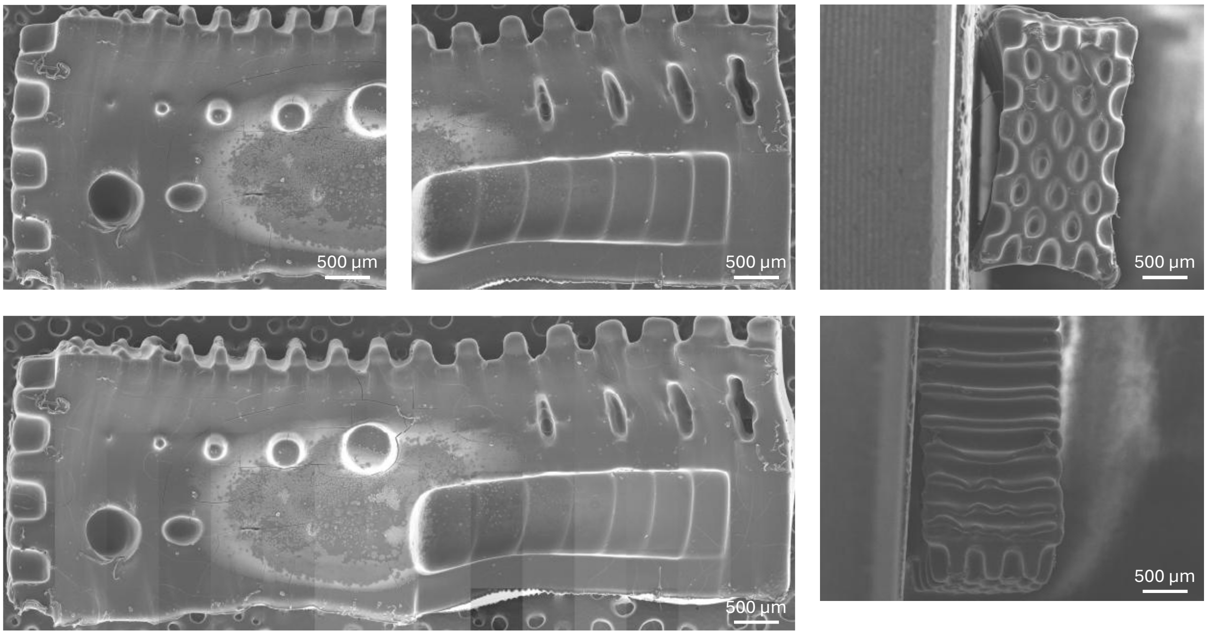


*Figure S12. SEM images of the benchmark (air dried). GelNB-GelSH63 10% (w/v) (0.05% (w/v) LAP) was VAM printed with set parameters: 56.26 mJ·cm$^{-2}$ dose and 0.3738 absorbance.*

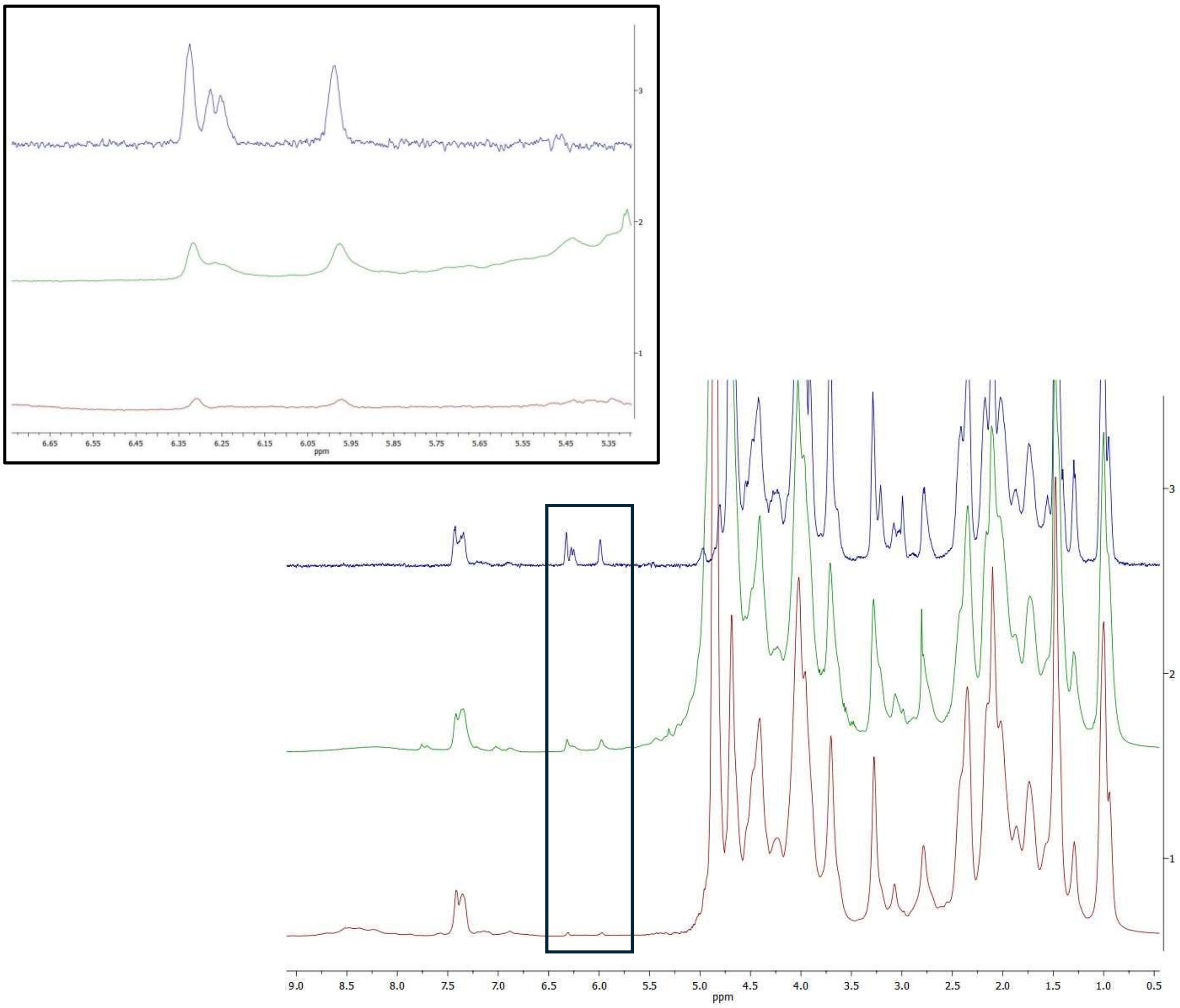

*Figure S13. [1]H-NMR and HR-MAS spectra of GelNB (blue, n°3), and the crosslinked materials prepared following the 2 methods for MSC encapsulation as described in the manuscript: film casted GelNB-GelSH (green, n°2) and VAM-printed GelNB-GelSH (red, n°1), corresponding with a double bond conversion of 41.7 and 95%, respectively.*

*Table S3. Overview of reference dose (obtained by rheology), refractive index (from refractometer measurements) and absorbance (from UVVIS) for varying % (v/v) of OptiPrep.*

| *OptiPrep [% (v/v)]* | *Reference dose [mJ•cm$^{-2}$]* | *Refractive index [-]* | *Absorbance [-]* |
|---|---|---|---|
| *0* | 33.9 | 1.34934 | 0.247 |
| *10* | N/A | 1.35771 | 0.181 |
| *15* | 35.2 | 1.36233 | 0.255 |
| *20* | 32.6 | 1.3661 | 0.288 |

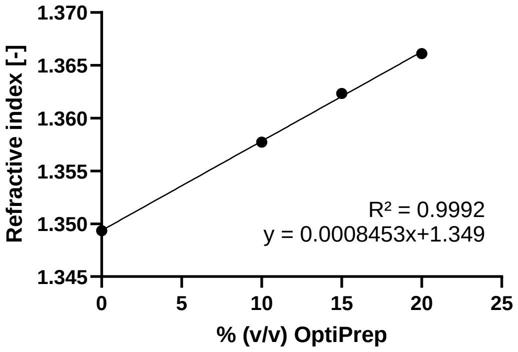


*Figure S14. Refractive index versus %(v/v) Optiprep, measured on bioresin composed of GelNB-GelSH63 10% (w/v) and MSCs (2·10$^{-6}$ cells per mL).*

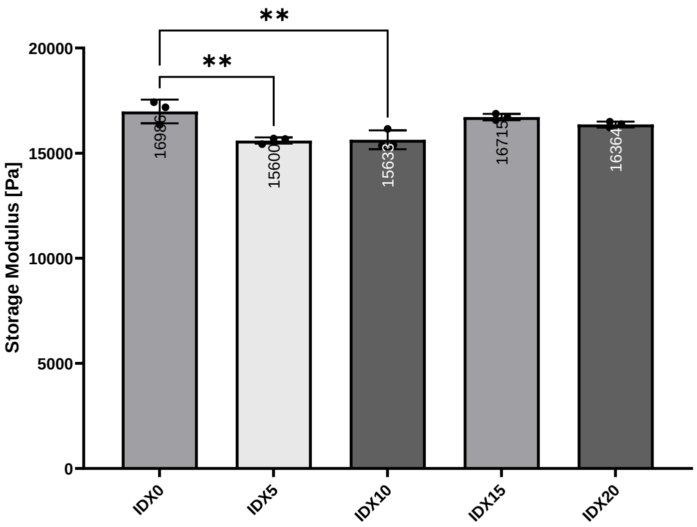


*Figure S15. Photorheological assessment of GelNB-GelSH63 photo-resins: storage (G') and loss (G") moduli as a function of OptiPrep concentration (% (v/v)). (* = p ≤ 0.05; ** = p ≤ 0.01; *** = p ≤ 0.001; **** = p ≤ 0.0001)*

### Photorheological Analysis of Bioresins with Varying OptiPrep Concentrations

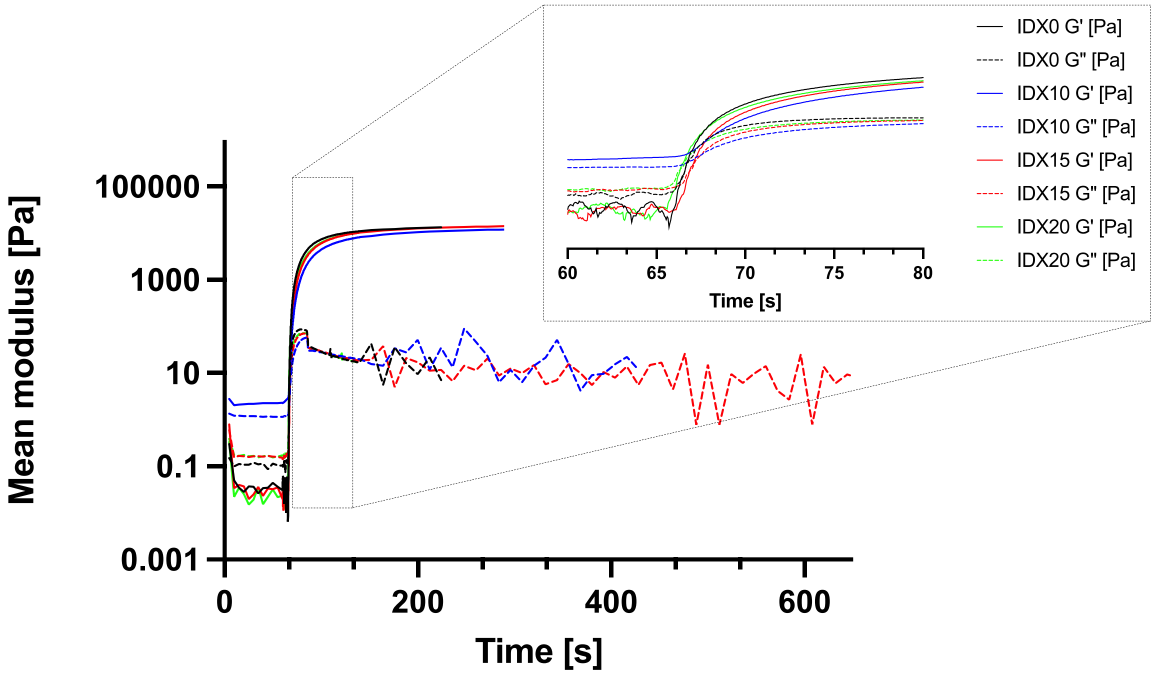


*Figure S16. Gel point determination using photorheology on GelNB-GelSH63 10% (w/v) and MSCs (2·10⁻⁶ cells per mL) on varying concentrations of iodixanol (IDX): 0, 10, 15 and 20 %(v/v) OptiPrep.*

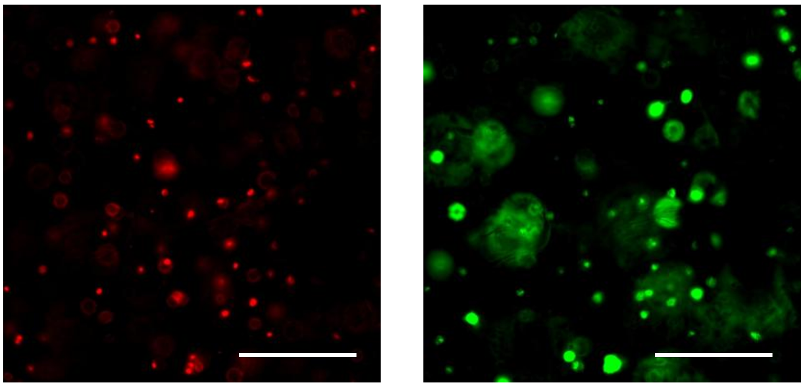

*Figure S17. Representative split-view live/dead staining image showing both viable (green, calcein AM-stained) and non-viable (red, propidium iodide-stained) cells (D0 of VAM-printed sample as example).*

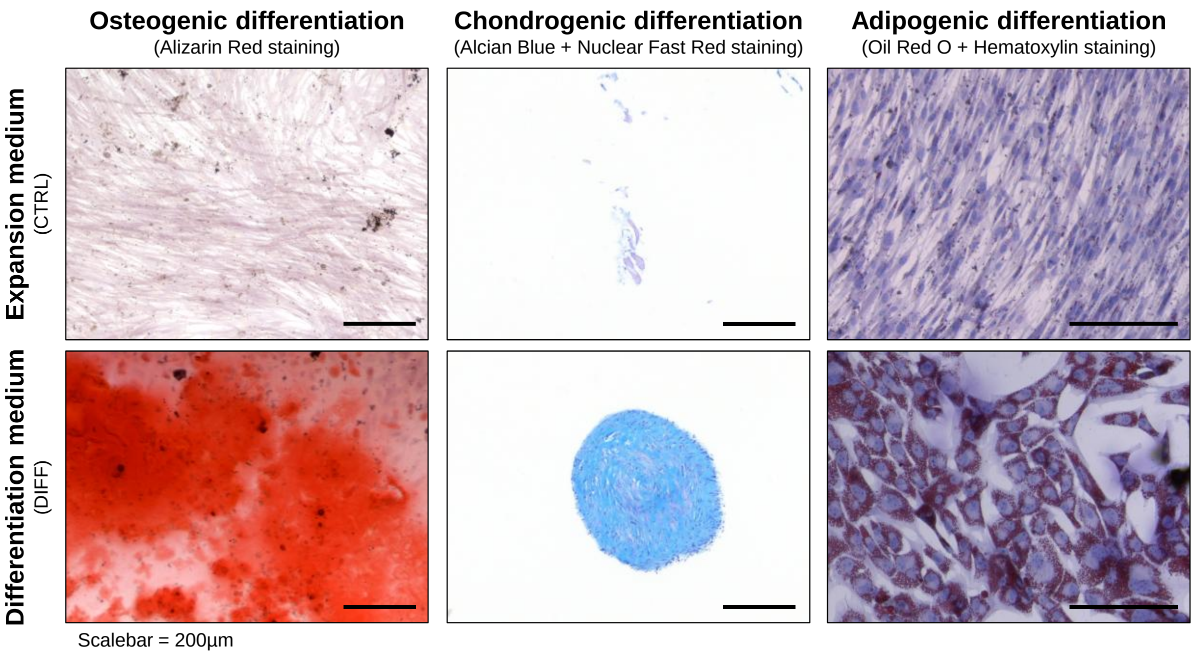


*Figure S18. Before encapsulating in film-cast of VAM-printed samples, MSCs were first differentiated to confirm their identity, according to the protocol of Heyman et al. (2022)* [26]
